# Stu2 has an essential kinetochore role independent of regulating microtubule dynamics

**DOI:** 10.1101/2022.09.09.507218

**Authors:** Ahmed Abouelghar, Joseph S. Carrier, Julia R. Torvi, Erin Jenson, Chloe Jones, Binnu Gangadharan, Elisabeth A. Geyer, Luke M. Rice, Brent Lagesse, Georjana Barnes, Matthew P. Miller

## Abstract

ch-TOG family proteins, including the budding yeast Stu2, are essential for spindle formation and chromosome segregation. Such functions depend on an array of activities ranging from microtubule nucleation, polymerization and depolymerization, to conferring tension sensitivity to kinetochores. This functional diversity makes it challenging to dissect these various functions and understand their relative importance. Here, we developed separation-of-function mutants and used artificial tethering tools to elucidate several important mechanistic insights into Stu2’s essential role. We show that Stu2’s microtubule polymerization activity depends on its basic linker region but is surprisingly dispensable for viability; that in fact, Stu2 carries out an essential kinetochore-associated function; and finally, that Stu2’s precise location within the kinetochore is critical for its function, suggesting a spatial separation mode of action may underlie its ability to confer tension sensitivity. Our findings highlight the significance of Stu2’s kinetochore role and provide insights into the molecular mechanisms by which it performs its various functions.

## INTRODUCTION

Cells use a microtubule cytoskeleton that provides shape and structure to the cell and facilitates the transport of organelles. This dynamic microtubule cytoskeleton is altered to accomplish various functions at different cell cycle stages. As the cell progresses into mitosis the cell must generate a mitotic spindle to accurately segregate chromosome pairs between the two newly forming cells. This process involves diverse microtubule functions including forming and orienting a bipolar mitotic spindle, establishing correct interactions between kinetochores and microtubules, and elongating the mitotic spindle to separate replicated sister chromatids during anaphase (Hagan and Hyams, 1988; Hayles et al., 1994; Kirschner and Mitchison, 1986).

The proper formation of the mitotic spindle requires the function of microtubule associated proteins (MAPs), which regulate the structure and function of the microtubule cytoskeleton. MAPs can perform diverse roles in regulating microtubules including nucleation, polymerization, depolymerization, and altering microtubule dynamics (reviewed in Gudimchuk & McIntosh, 2021). They also act to link microtubules with other cellular elements, such as organelles, and regulate other MAPs by directing cellular localization (Bodakuntla et al., 2019).

One such family of MAPs is the ch-TOG family of proteins, which conduct conserved functions across all eukaryotes (Al-Bassam and Chang, 2011; Kinoshita et al., 2002; Ohkura et al., 2001). These proteins, including the budding yeast ortholog Stu2, are best known for their well-studied function in stimulating microtubule polymerization (Ayaz et al., 2014; Brouhard et al., 2008; Fox et al., 2014; Geyer et al., 2018; Nithianantham et al., 2018); however, ch-TOG proteins have additional roles in the cell. For example, ch-TOG proteins regulate the dynamics and disassembly of microtubules (Humphrey et al., 2018; Kosco et al., 2001; Podolski et al., 2014; Usui et al., 2003; Van Breugel et al., 2003). They also nucleate and anchor microtubules at the microtubule organizing centers (Flor-Parra et al., 2018; Gunzelmann et al., 2018; King et al., 2020; Thawani et al., 2018). At the kinetochore, ch-TOG proteins aid in kinetochore capture by the spindle microtubules (Gandhi et al., 2011; Kitamura et al., 2010). Furthermore, this family of proteins is implicated in correcting erroneous kinetochore-microtubule attachments. Stu2 confers tension sensitivity to purified kinetochores in vitro, stabilizing high-tension attachments while destabilizing low-tension ones (Miller et al., 2016). Given the multitude of functions these proteins fulfill, it is not surprising that *stu2* mutants result in chromosome segregation defects. Nevertheless, it remains unclear to what extent these proteins directly contribute to establishing proper kinetochore-microtubule attachments and whether this function is required for cell viability.

To perform these diverse functions, ch-TOG proteins utilize various domains to interact with microtubules and localize to distinct subcellular locations. Among these are conserved TOG domains which bind tubulin heterodimers at or near microtubule plus ends, where they stimulate large changes in the addition or loss of tubulin subunits to the microtubule polymer (Al-Bassam et al., 2006; Ayaz et al., 2014; Brouhard et al., 2008; Geyer et al., 2018; Nithianantham et al., 2018; Podolski et al., 2014; Van Breugel et al., 2003; Widlund et al., 2011). They also possess a carboxy-terminal binding segment (CTS), well-known for facilitating numerous binding interactions and directing them to various cellular locations (Gunzelmann et al., 2018; Stangier et al., 2018; Thawani et al., 2018; Zahm et al., 2021). Finally, ch-TOG proteins engage with the negatively charged microtubules through a highly basic linker region. Importantly, this ‘basic linker’ plays a crucial role in various functions and is implicated in nearly all described activities of the ch-TOG family. These functions include facilitating their microtubule lattice diffusion (Al-Bassam et al., 2006; Geyer et al., 2018; Wang and Huffaker, 1997; Widlund et al., 2011), oligomerizing the γ-TuSC in conjunction with the γ-tubulin receptor Spc72 to facilitate microtubule nucleation (Gunzelmann et al., 2018), and contributing to the kinetochore function of the human ortholog, ch-TOG (Herman et al., 2020).

Considering the observation that the basic linker domain is implicated in many, if not all, ch-TOG family members functions, one would expect this domain to be essential. However, our previous work examining the yeast family member Stu2 indicates that a specific patch comprising 15 conserved residues within the entire 97 amino acid basic linker domain is both necessary and sufficient for viability (Herman et al., 2020). Given the improbability of this 15-residue patch alone performing all the functions attributed to the entire basic linker, this observation raises questions about the essential cellular role of Stu2, including its most well-known function as a microtubule polymerase.

In this study, we show that, as previously proposed, Stu2’s basic linker is crucial for its microtubule polymerization and depolymerization activity. Furthermore, we reveal that the conserved patch within Stu2’s basic linker comprises Stu2’s nuclear localization signal. Surprisingly, localizing Stu2 to the nucleus is the only activity of the basic linker required for cell viability, indicating that Stu2’s ability to induce large changes in microtubule dynamics is dispensable. Building from these findings, we engineered a lethal *stu2* mutant that lacks the ability to properly interact with many of Stu2’s binding partners, preventing it from localizing to its subcellular sites of action. Cell viability is restored when this lethal mutant is artificially tethered near its native binding site at the C-terminus of Nuf2, a component of the outer kinetochore Ndc80 complex, but not when tethered to other known Stu2 interactors. This demonstrates that Stu2 plays an essential role at the kinetochore. To investigate this kinetochore function, we examined a model we previously proposed for how Stu2 might confer tension sensitivity to kinetochores (Miller et al., 2016). We proposed that at low tension, Stu2 destabilizes attachments by competing with (or binding to and inhibiting) the microtubule binding of another element at the kinetochore, such as the CH domains of the Ndc80 complex or the Dam1 complex. This destabilization activity is suppressed by spatial separation under high tension as the outer kinetochore structure stretches. With the genetic tools we developed, we tested this model by modulating Stu2’s precise position within the outer kinetochore. Remarkably, we found that artificially localizing Stu2 near the CH domains of the Ndc80 complex led to cell death and an accumulation of unattached kinetochores. These findings support a spatial separation model, where high tension separates Stu2 from other microtubule-binding factors, stabilizing the attachments. Overall, our results demonstrate that Stu2’s kinetochore function is essential for proper mitotic progression and cell viability, independent of its role in regulating microtubule dynamics, and provide insight into a plausible mechanism of action. These findings significantly impact our understanding of how ch-TOG proteins function to ensure proper mitotic progression.

## RESULTS

### Two positive residues within a conserved patch of amino acids in Stu2’s basic linker region are essential for cell viability

Our earlier research revealed that a distinct patch in the basic linker of Stu2 consisting of 15 conserved residues (Stu2^592-606^) is not only necessary but also sufficient for cell viability (Herman et al., 2020; Fig. S1A-C). As this patch constitutes only a small portion of the entire basic linker domain (i.e. 15 of 97 residues), we hypothesized that it is unlikely that it alone can carry out all of the basic linker functions. This idea motivated us to investigate the function of the conserved patch. To examine which elements of this conserved patch are required for cellular viability, we made mutants spanning this region and performed a serial dilution spot viability assay. Strains contained an endogenously expressed *STU2* allele tagged with an auxin-inducible degron (i.e. *stu2-AID*), allowing for degradation of this copy of Stu2 in the presence of auxin (Fig. S1D), and expressed mutant variants of *stu2* from an ectopic locus (under control of the native *STU2* promoter). We mutated each of the 15 amino acids in the conserved patch to alanine and found that two highly conserved positive residues, Stu2^K598^ and Stu2^R599^, are required for cell viability. Alanine mutations of Stu2^K598^ and Stu2^R599^ either independently, or together (henceforth referred to as *stu2^KR/AA^*) cause severe viability defects (Fig. 1A-C and Fig. S1E-I). Furthermore, cells expressing *stu2^KR/AA^*display pleiotropic mitotic defects, including a short and misoriented mitotic spindle (*stu2^KR/AA^* 29% misoriented vs *STU2* 17%), as well as defects in bi-lobed kinetochore distribution (*stu2^KR/AA^*28% mono-lobed vs *STU2* 1%) and the presence of unattached kinetochores (*stu2^KR/AA^* 40% vs *STU2* 2%; Fig. 1D and Fig. S1J-K). These pleiotropic defects observed in *stu2^KR/AA^*-expressing cells suggest that K598 and R599 play a crucial role in multiple of Stu2’s functions.

**Figure 1.**
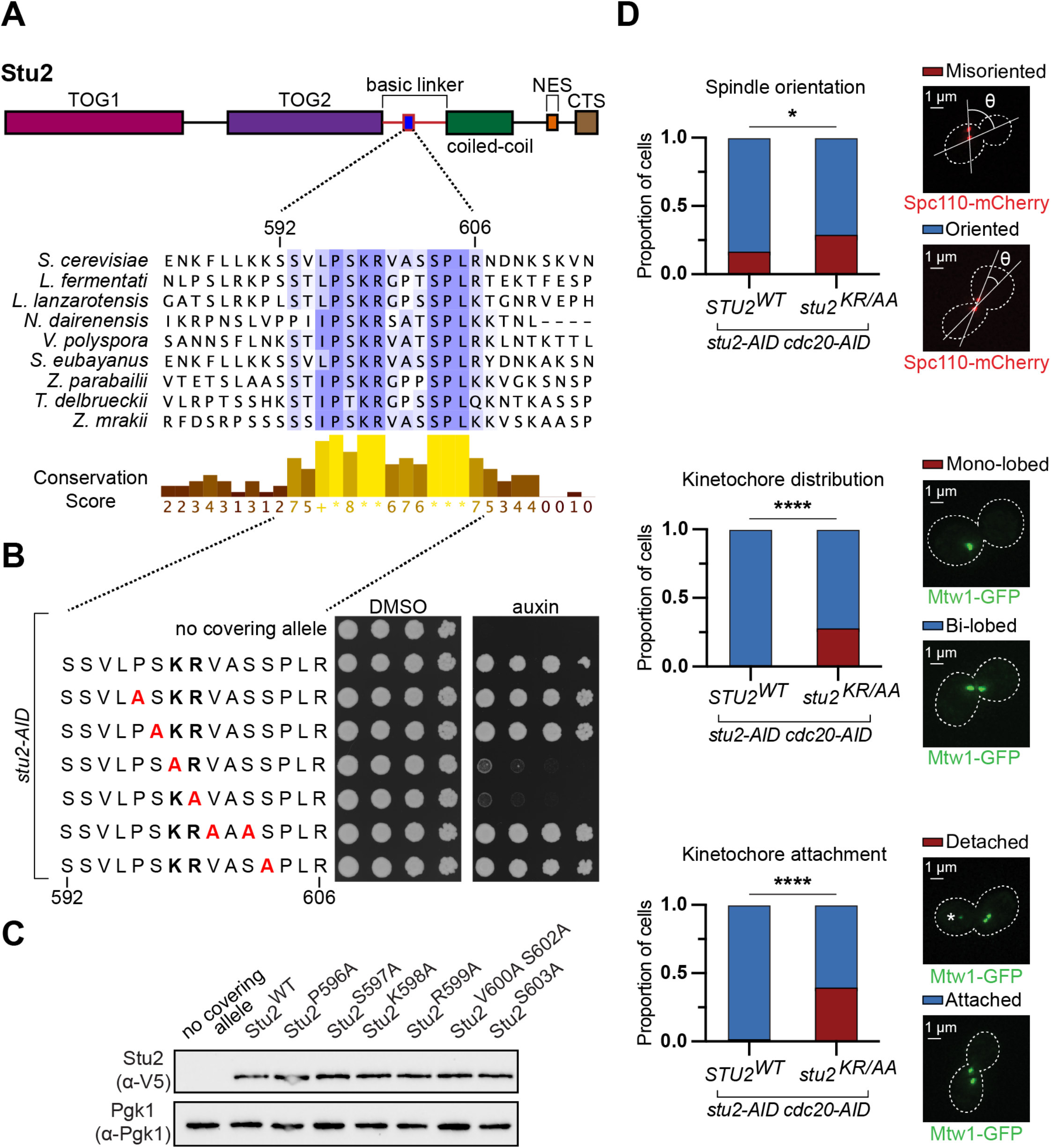
Two positive residues within a conserved patch in Stu2’s basic linker region are essential for cell viability. (A) Top: A schematic of Stu2’s domains (NES, nuclear export sequence; CTS, c-terminal segment). Bottom: Multiple sequence alignment across 9 different fungal species reveals a conserved patch of amino acids within Stu2’s basic linker region. (B) Cell viability was analyzed in *stu2-AID* strains without a covering allele (M619) or ectopically expressing *stu2* alleles (*STU2^WT^*, M622; *stu2^P596A^*, M1290; *stu2^S597A^*, M1299; s*tu2^K598A^*, M954; *stu2^R599A^*, M956; *stu2^V600A^ ^S602A^*, M1383; or *stu2^S603A^*, M802) by plating five-fold serial dilutions in the presence of DMSO (left) or auxin (right) to degrade the endogenous Stu2-AID protein. (C) Exponentially growing *stu2-AID* strains without a covering allele (M619) or ectopically expressing *stu2-3V5* alleles (*STU2^WT^*, M622; *stu2^P596A^*, M1290; *stu2^S597A^*, M1299; s*tu2^K598A^*, M954; *stu2^R599A^*, M956; *stu2^V600A^ ^S602A^*, M1383; or *stu2^S603A^*, M802) were treated with auxin 30 min prior to harvesting. Protein lysates were subsequently prepared and expression of Stu2- 3V5 proteins were analyzed by immunoblotting. Pgk1 is shown as a loading control. (D) Exponentially growing *stu2-AID cdc20-AID* cells ectopically expressing *stu2* alleles (*STU2^WT^*, M1914; or *stu2^KR/AA^*, M1922) were treated with auxin for 2 h. Cells were fixed and both the Spc110-mCherry signal and Mtw1-GFP signal were imaged. Top: Cells with a mitotic spindle oriented within 45 degrees of the mother-daughter bud axis versus greater than 45 degrees were counted. Proportion of cells with a spindle oriented greater than 45 degrees relative to the mother-daughter bud axis, n=101-103 (left). Representative cells with an oriented versus misoriented spindle (right). Middle: Cells with a mono-lobed GFP puncta versus bi-lobed GFP punctae were counted. Proportion of cells for each strain with a mono-lobed GFP puncta are shown, n=101-103 cells (left). Representative images of cells with a mono-lobed versus bi-lobed GFP puncta (right). Bottom: Cells with a GFP puncta isolated from the normal bipolar GFP punctae were counted and percentage of cells with an isolated puncta for each strain are shown, n=101-103 cells (right). Representative cells with and without an isolated GFP puncta (left). The GFP puncta is marked with a *. P values were determined using a Fisher’s exact test (* = p<0.05, **** = p<0.0001).

### Conserved patch constitutes Stu2’s NLS

We postulated that one plausible explanation for the diverse mitotic defects observed in *stu2^KR/AA^*mutants could be alterations in Stu2’s capacity to accurately localize to the nucleus during mitosis. To test this hypothesis, we fluorescently labeled the nuclear envelope by tagging the nucleoporin protein, Nup2, with mKate2 and compared the ratio of nuclear to cytoplasmic Stu2-GFP intensities in metaphase-arrested cells expressing Stu2^WT^ vs Stu2^KR/AA^. We observed substantially less nuclear localization in cells expressing Stu2^KR/AA^ (nuclear to cytoplasmic GFP ratio: 0.59 ± 0.26) compared to Stu2^WT^ (1.57 ± 0.35), suggesting significantly disrupted nuclear localization of the Stu2^KR/AA^ mutant (Fig. 2A). To further test the role of Stu2^K598^ ^R599^ in localizing Stu2 to the nucleus, we examined the ability of Stu2^KR/AA^ to interact with the α-subunit of the importin complex, Kap60, which binds directly to the NLS of nuclear localization sequence-containing proteins (Christie et al., 2016; Enenkel et al., 1995). We tagged Kap60 with a Flag epitope and used α-Flag conjugated magnetic beads to pull down Kap60 and examine binding of V5-tagged Stu2^WT^ or Stu2^KR/AA^. Here we observed that the co-immunopurified Stu2^KR/AA^ signal was significantly reduced compared to the signal for Stu2^WT^ (Fig. 2B). These observations are consistent with the hypothesis that the conserved patch constitutes a portion of the nuclear localization sequence (NLS) of Stu2. This idea is also consistent with work from Kosugi *et al*. who defined a nuclear localization sequence class with the consensus sequence (P/R)XXKR(^DE)(K/R), where (^DE) indicates any residue except for D/E (Kosugi et al., 2009).

**Figure 2.**
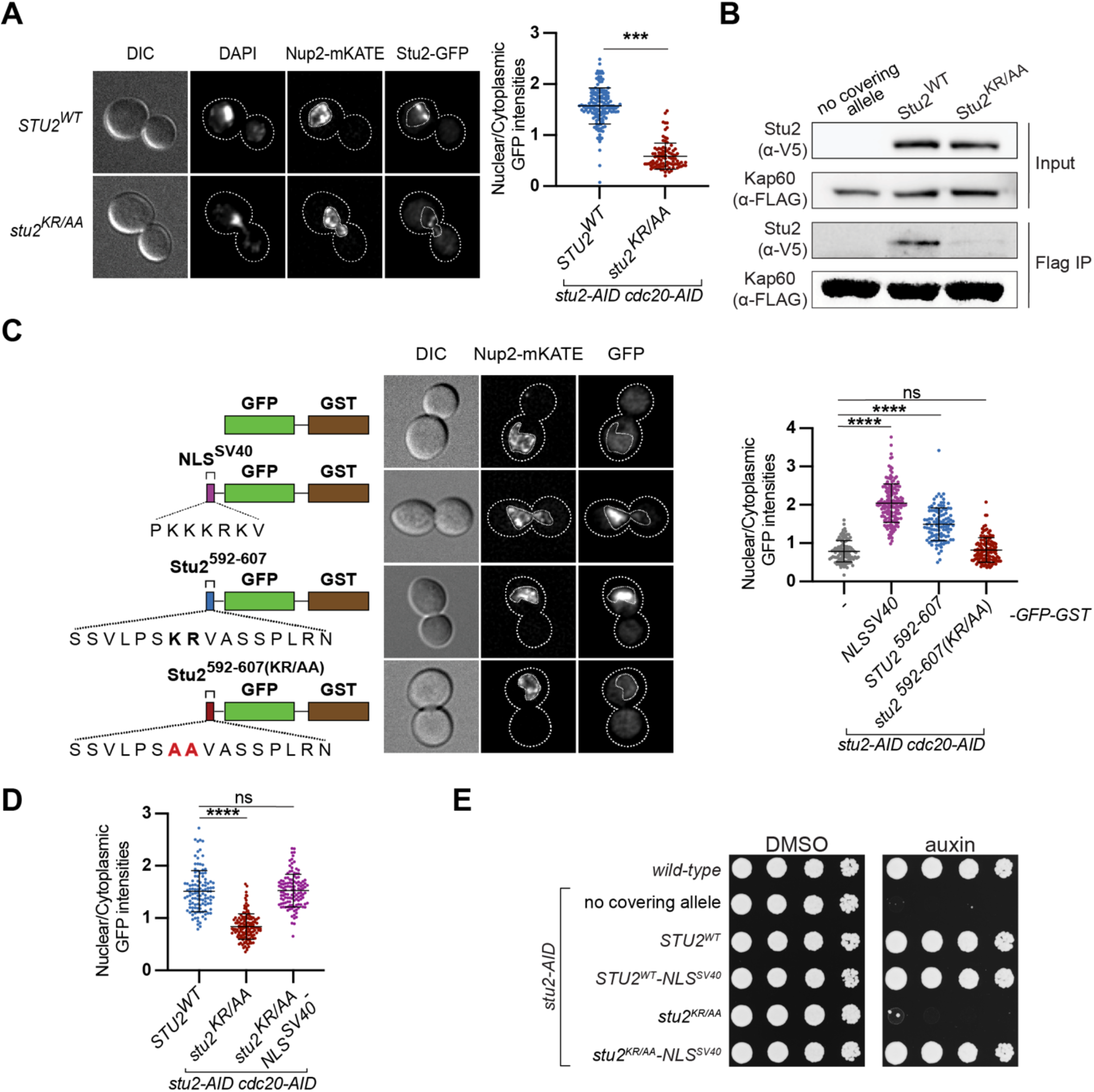
Stu2^592-607^ is required to import Stu2 into the nucleus. A) Exponentially growing *NUP2-mKATE stu2-AID cdc20-AID* cells ectopically expressing *stu2- GFP* alleles (*STU2^WT^*, M2208; or *stu2^KR/AA^*, M2210) were treated with auxin for 2 h to arrest in metaphase. Cells were fixed and imaged for DAPI, Nup2-mKATE, and Stu2-GFP. Left: Representative images of DIC, DAPI, Nup2-mKATE, and Stu2^WT^-GFP (top) or Stu2^KR/AA^-GFP signals (bottom) with a representative outline of the nucleus. Right: Quantification of the ratio of the nuclear to cytoplasmic GFP signal intensities with each data point representing a measurement from a single cell. Mean and standard deviation are shown. n=96-160 cells; P values were determined using a two-tailed unpaired *t* test (*** = p<0.001). (B) Exponentially growing *stu2-AID* cells expressing Kap60-3FLAG without a covering *STU2* allele (M2350) or ectopically expressing *stu2-3V5* alleles (*STU2^WT^*, M2435; or *stu2^KR/AA^*, M2436) were treated with auxin 30 min prior to harvesting. Protein lysates were subsequently prepared; Kap60 and associated binding partners were purified by α-Flag immunoprecipitation (IP), and associated Stu2-3V5 proteins were analyzed by immunoblotting. (C) Exponentially growing *stu2-AID cdc20-AID* cells ectopically expressing *GFP-GST* fused constructs (*GFP-GST*, M2390; *NLS^SV40^-GFP-GST*, M2391; *stu2^592-607^-GFP-GST*, M2392; or *stu2^592-607(KR/AA)^-GFP-GST*, M2437) were treated with auxin for 2 h. Cells were fixed and the Nup2-mKATE and GFP signals were imaged. Left: Schematic of the constructs used. Middle: Representative images of DIC, Nup2-mKATE, and GFP signals for each strain with an outlined nucleus. Right: Quantification of the ratio of the nuclear to cytoplasmic GFP signal intensities. Each data point represents this ratio for a single cell. The nuclear pool of GFP-GST construct suggests that some GFP-GST can freely diffuse into the nucleus. Mean and standard deviation are shown. n=111-161 cells; P values were determined using an ordinary one-way ANOVA followed by a Tukey’s multiple comparisons test (**** = p<0.0001). (D) Exponentially growing *stu2-AID cdc20-AID* cells ectopically expressing *stu2-GFP* alleles (*STU2^WT^*, M2208; *stu2^KR/AA^*, M2210; or *stu2^KR/AA^-NLS^SV40^*, M2353) were treated with auxin for 2 h to arrest in metaphase. Cells were fixed and Nup2-mKATE, and Stu2-GFP signals were imaged. The ratio of the nuclear to cytoplasmic GFP signal intensities was quantified. Each data point represents this ratio for a single cell. Mean and standard deviation are shown. n=114-137 cells; P values were determined using an ordinary one-way ANOVA followed by a Tukey’s multiple comparisons test (**** = p<0.0001). (E) Cell viability was analyzed in a wild-type strain (M3) and *stu2-AID* strains without a covering allele (M619) or ectopically expressing *stu2* alleles (*STU2^WT^*, M2103; *STU2^WT^-NLS^SV40^*, M2105; *stu2^KR/AA^*, M2225; or *stu2^KR/AA^-NLS^SV40^*, M2199) by plating five-fold serial dilutions in the presence of DMSO (left) or auxin (right) to degrade the endogenous Stu2-AID protein.

Since Stu2^K598^ ^R599^ are necessary for nuclear localization and Kap60 binding, we next wanted to assess if Stu2^592-607^ is sufficient to promote nuclear localization. We engineered a fusion protein containing Stu2’s conserved patch fused to GFP-GST ectopically expressed under the *STU2* promoter. The GST protein was included to make the protein construct large enough to limit the amount of GFP freely diffusing into the nucleus (Okada et al., 2014). When examined, GFP-GST alone primarily localized to the cytoplasm in metaphase-arrested cells (nuclear to cytoplasmic GFP signal ratio of 0.79 ± 0.28). As expected, the addition of the NLS of the SV40 virus large T-antigen protein, NLS^SV40^, promoted GFP nuclear import (nuclear to cytoplasmic GFP signal ratio of 2.05 ± 0.50). Importantly, fusion of Stu2^592-607^ to GFP-GST was sufficient to promote nuclear import, and this activity was disrupted when the Stu2^K598^ ^R599^ residues were mutated in this minimal context (nuclear to cytoplasmic ratio of GFP signal of 1.49 ± 0.43 for Stu2^592-607^-GFP-GST and 0.82 ± 0.32 for Stu2^592-607(KR/AA)^-GFP-GST; Fig. 2C and Fig. S2A). These results demonstrate that the conserved patch located within Stu2’s basic linker constitutes Stu2’s NLS.

Finally, to test if the decreased nuclear localization was responsible for the viability defects of *stu2^KR/AA^* mutants, we examined if cell viability was rescued by increasing the nuclear accumulation of Stu2^KR/AA^. We tested this idea in two ways: First, we fused NLS^SV40^ to Stu2^KR/AA^ and found that both the growth and nuclear localization defects were completely rescued when fused with NLS^SV40^ (Fig. 2D-E). Secondly, we modulated Stu2’s nuclear export by the previously described phospho-regulation of its nuclear export signal and observed similar results (van der Vaart et al., 2017; Fig S2B). Together, these results indicate that Stu2’s nuclear localization is essential for cell viability, and that the predominant function of Stu2^K598^ ^R599^ is to promote Stu2’s import into the nucleus.

### Directing Stu2 nuclear localization is the only essential function of Stu2’s entire basic linker region

The basic linker is thought to be important for Stu2’s ability to regulate microtubule assembly and disassembly. Considering that only a small portion of the basic linker, Stu2^592-607^, is both necessary and sufficient for maintaining cell viability (Fig. S1B and Fig. S3A), this raised the possibility that the only essential function of Stu2’s entire basic linker is to facilitate nuclear localization. To explore this idea, we examined whether we could restore the viability phenotype of a basic linker deletion (Stu2^ΔBL^) by fusing an NLS^SV40^. To our surprise, C-terminal fusion of NLS^SV40^ completely restored the viability of the entire basic linker deletion (Fig. 3A and Fig. S3B). Considering that the number of positive charges within the basic linker was previously shown to be crucial for Stu2’s ability to control microtubule dynamics in vitro, it is quite surprising that the basic linker is dispensable for cell viability (Geyer et al., 2018). There are two potential explanations for this phenomenon: either Stu2 can regulate microtubule dynamics independently of the basic linker, or Stu2’s role in controlling microtubule dynamics is not essential.

**Figure 3.**
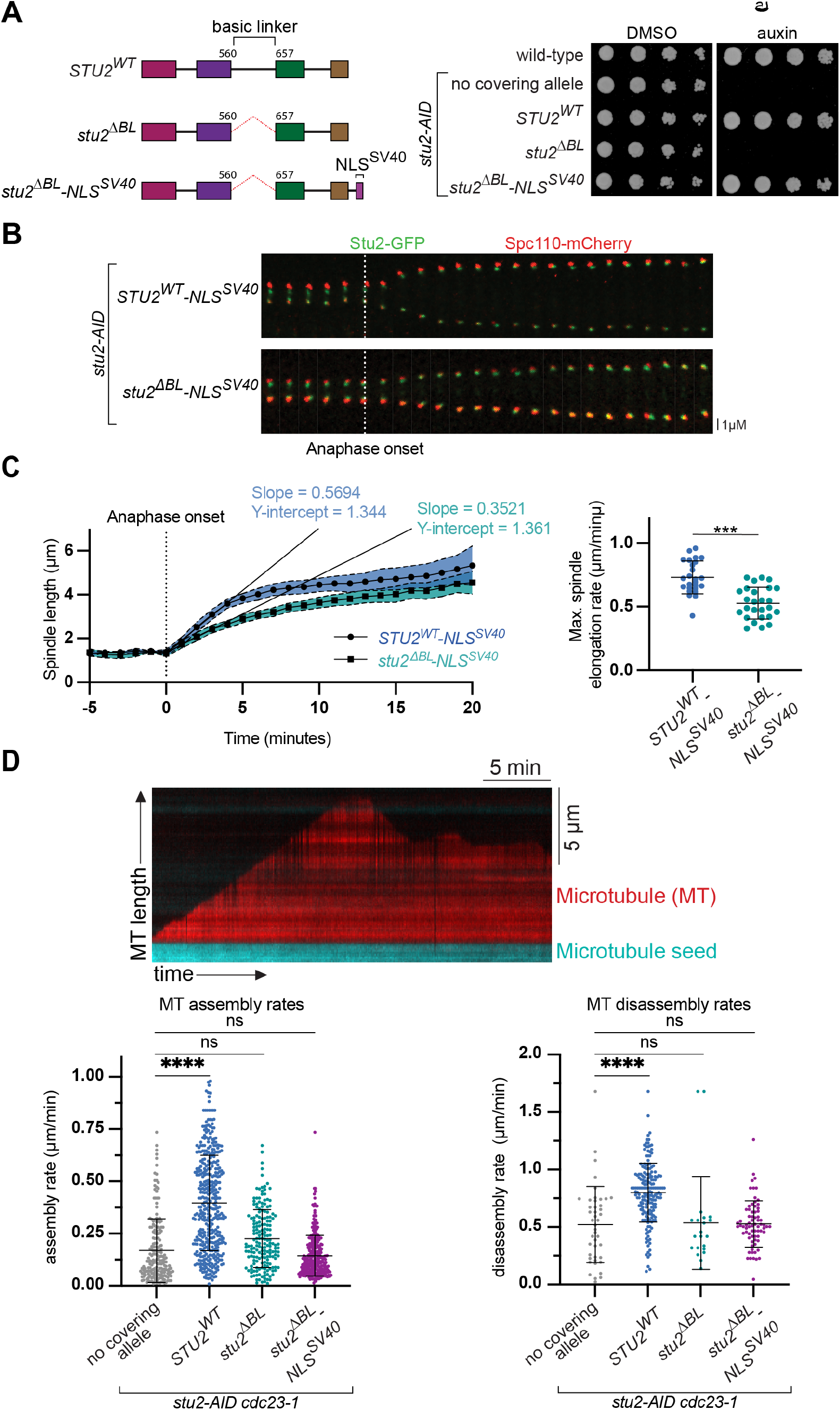
Stu2’s basic linker region is dispensable for cell viability outside of localizing Stu2 to the nucleus. (A) Left: A schematic of Stu2 mutants. Right: Cell viability was analyzed in a wild-type strain (M3) and *stu2-AID* strains without a covering allele (M619) or ectopically expressing *stu2* alleles (*STU2^WT^*, M622; *stu2^ΔBL^*, M973; or *stu2^ΔBL^-NLS^SV40^*, M3722) by plating five-fold serial dilutions in the presence of DMSO (left) or auxin (right) to degrade the endogenous Stu2-AID protein. (B) Exponentially growing *stu2-AID* cells ectopically expressing *stu2-GFP* (*STU2^WT^-NLS^SV40^*, M2729; *stu2^ΔBL^-NLS^SV40^*, M2731) were treated with auxin for 30 min to degrade Stu2-AID. Spc110-mCherry and Stu2-GFP signals were imaged every min. Representative time course of a cell progressing through anaphase beginning 5 min prior to anaphase onset until 20 min post-anaphase onset (merge of Spc110-mCherry and Stu2-GFP signals shown). The white dotted line indicates anaphase onset. (C) Left: Mean and 95% confidence intervals of spindle length are shown from 5 min prior to anaphase onset to 20 min post-anaphase onset. The black dotted line indicates anaphase onset. Lines with indicated slopes and y-intercepts are lines of best fit during the period of maximum rate of spindle elongation. n=25-26. Right: Calculated maximum rates of spindle elongation over a 2-min period for each individual cell. Each data point represents a single cell. Mean and standard deviation are shown. n=25-26; P values were determined using a two-tailed unpaired *t* test (*** = p<0.001). (D) Exponentially growing *GFP-TUB1 stu2-AID cdc23-1* cells with no covering allele (M2166) or ectopically expressing *stu2-Halo* (*STU2^WT^*, M2231; *stu2^ΔBL^*, or M2360; *stu2^ΔBL^-NLS^SV40^*, M3516) were shifted to a non-permissive temperature for 3 h to arrest cells in metaphase and treated with auxin 30 min before harvesting to degrade Stu2-AID. Whole cell lysate was incubated with microtubule seeds, and growth and shrinkage rates of microtubule extensions were determined. Top: Representative kymograph of *STU2^WT^*(red, microtubule extension; cyan, seed). Rates of microtubule assembly or disassembly are determined by the slope of microtubule extension growth and shrinkage over time. Bottom left: Assembly rates of microtubules. Each data point represents an independent microtubule assembly event. Mean and standard deviation are shown. n=182-397 growth events. Right: Disassembly rates of microtubules. Each data point represents an independent microtubule disassembly event. Mean and standard deviation are shown. n=21-179 shrinkage events; P values were determined using an ordinary one-way ANOVA followed by a Tukey’s multiple comparisons test (**** = p<0.0001).

To test if the basic linker is required for regulating microtubule dynamics, we used two methods: spindle elongation in cells and microtubule dynamics in vitro. First, we examined anaphase spindle elongation rates as a readout of microtubule polymerization activity (Severin et al., 2001). We found that the rate at which Stu2^ΔBL^-NLS^SV40^ cells elongated their spindle during anaphase was significantly reduced compared to cells expressing Stu2^WT^-NLS^SV40^ (Fig. 3B-C), consistent with the basic linker being required for microtubule polymerization in vivo. Next, we examined whether the basic linker is required to regulate microtubule assembly and disassembly in vitro. To achieve this, we adapted a previously established in vitro whole cell lysate assay (Bergman et al., 2019) to investigate the influence of various Stu2 variants on microtubule dynamics. As expected, we found that lysate containing Stu2^WT^ compared to lysate depleted of Stu2 displayed a significant increase in the microtubule assembly rate (Stu2^WT^, 0.40 ± 0.23 µm/min; no covering allele, 0.17 ± 0.15 µm/min). Consistent with the idea that the basic linker domain is important in microtubule polymerase function (Geyer et al., 2018; Widlund et al., 2011), both Stu2^ΔBL^ and Stu2^ΔBL^-NLS^SV40^-containing lysates had substantially decreased rates of microtubule elongation (Stu2^ΔBL^, 0.23 ± 0.14 µm/min; Stu2^ΔBL^-NLS^SV40^, 0.14 ± 0.10 µm/min; Fig. 3D). Furthermore, both Stu2^ΔBL^ and Stu2^ΔBL^-NLS^SV40^ were also defective in microtubule depolymerization activity and their ability to induce microtubule catastrophe (Fig. 3D and Fig. S3C-K).

These findings demonstrate that Stu2’s basic linker is required for its ability to induces changes in microtubule dynamics. We note a strong lack of correlation between Stu2 variants’ capacity to stimulate microtubule assembly and disassembly and their ability to support cell viability (compare Fig. 3D and 3A, respectively). This observation strongly indicates that Stu2’s ability to induce substantial alterations in mitotic spindle dynamics is unexpectedly dispensable for cell viability. Moreover, these findings suggest that Stu2^ΔBL^-NLS^SV40^ effectively separates Stu2’s essential activity from its microtubule regulatory functions. Consequently, this variant serves as a crucial tool for elucidating Stu2’s essential function.

### Stu2’s kinetochore activity is essential for cell viability

To gain insight into what essential function Stu2 carries out in the cell, we wanted to determine the subcellular location where this activity is required. Our overall approach involved 1) disrupting Stu2’s localization by perturbing its binding to interacting proteins; and 2) artificially restoring its localization to specific sites and examining whether this ‘re-tethering’ was sufficient to restore cellular function. Stu2 contains a carboxy-terminal binding segment (CTS) that has been well-characterized as facilitating many of Stu2’s binding interactions, including interactions with the yeast kinetochore through the Ndc80 complex (Miller et al., 2019; Zahm et al., 2021), with the cytoplasmic face of the yeast spindle pole body through Spc72 (Chen et al., 1998; Gunzelmann et al., 2018; Usui et al., 2003; Wang and Huffaker, 1997) and with the MAPs Bim1 and Bik1 to regulate microtubule dynamics (Wolyniak et al., 2006). Thus, we wanted to test if loss of the CTS in the sensitized basic linker deletion mutant would be lethal. Indeed, this was the case, as *stu2^ΔBL ΔCTS^-NLS^SV40^* mutants are completely inviable (Fig 4A and S4A). As a control we combined a basic linker deletion with a deletion of a different domain of Stu2, its homodimerization coiled-coil domain. The *stu2^ΔBL^ ^ΔCC^-NLS^SV40^* mutant exhibited only a subtle growth defect compared to the coiled-coil deletion, *stu2 ^ΔCC^-NLS^SV40^*. These findings indicate that the Stu2^ΔBL^ ^ΔCTS^-NLS^SV40^ variant is unable to fulfill Stu2’s essential function, which we hypothesize is due to the disruption of its necessary cellular localization (also see Supplementary Discussion Point 1).

**Figure 4.**
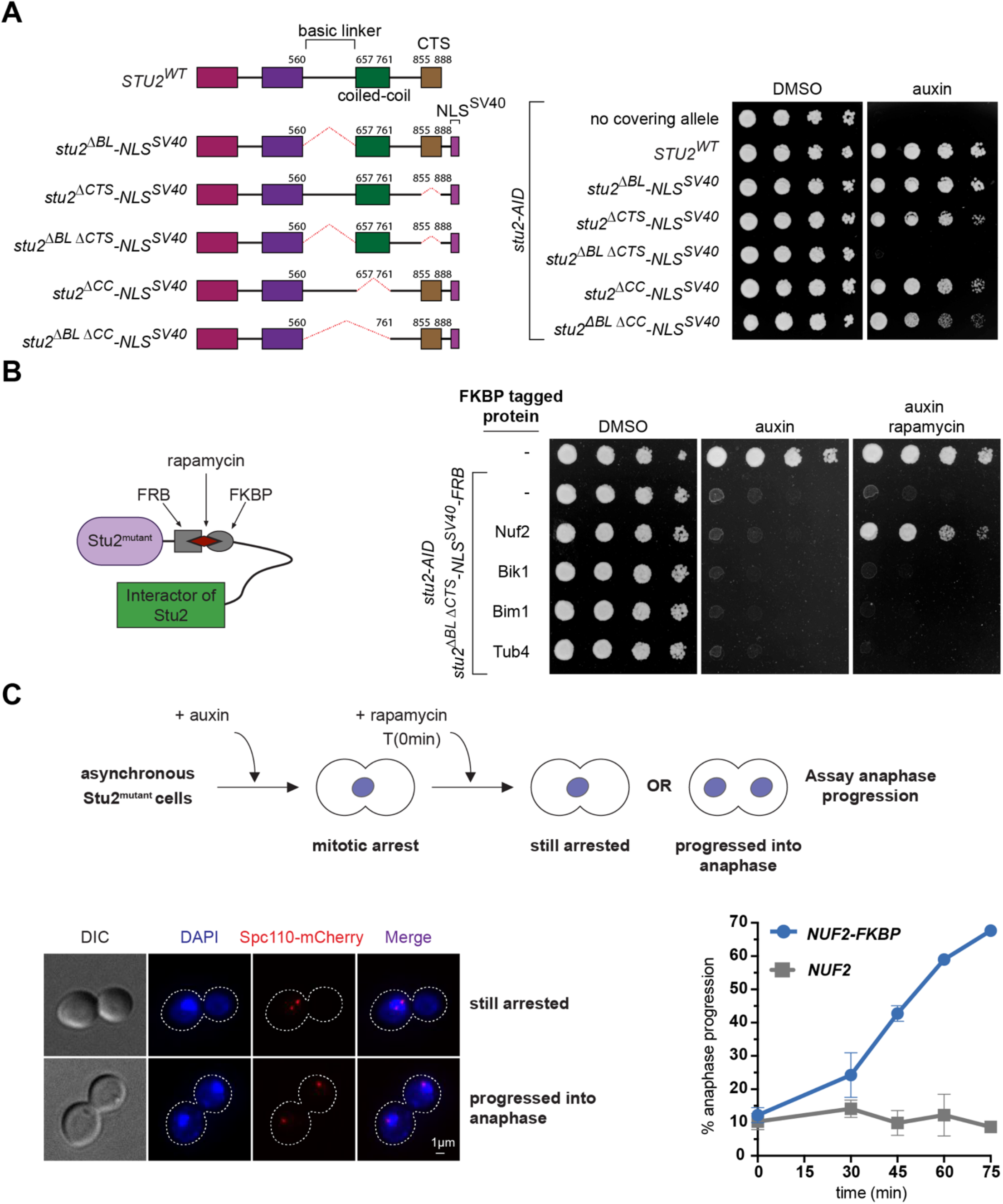
Stu2’s kinetochore activity is required for cell cycle progression. (A) Left: A schematic of Stu2 mutants. Right: Cell viability was analyzed in *stu2-AID* strains without a covering allele (M619) or ectopically expressing *stu2* alleles (*STU2^WT^*, M622; *stu2^ΔBL^- NLS^SV40^*, M3722; *stu2^ΔCTS^-NLS^SV40^*, M3942; *stu2^ΔBL^ ^ΔCTS^-NLS^SV40^*, M3727; *stu2^ΔCC^-NLS^SV40^*, M3940; or *stu2^ΔCC^ ^ΔCTS^-NLS^SV40^*, M3729) by plating five-fold serial dilutions in the presence of DMSO (left) or auxin (right) to degrade the endogenous Stu2-AID protein. (B) Left: A schematic illustrating rapamycin-induced dimerization of FRB and FKBP, demonstrating how this system was used to artificially tether Stu2 mutants to various Stu2 interactors. Right: Cell viability was analyzed in a *TOR1-1 fpr1Δ* strain that is otherwise wild- type (M1375) and *TOR1-1 fpr1Δ stu2-AID* strains with *stu2^ΔBL^ ^ΔCTS^-NLS^SV40^-FRB* covering allele and without any FKBP tagged gene (M4515) or endogenously expressing FKBP tagged genes (*NUF2*, M4366; *BIK1*, M4369; *BIM1*, M4371; or *TUB4*, M4367) by plating five-fold serial dilutions in the presence of DMSO (left), auxin (middle) to degrade the endogenous Stu2-AID protein, or auxin and rapamycin (right) to induce the dimerization of FRB and FKBP and degrade the endogenous Stu2-AID protein. (C) Top: A schematic of the experimental design. Strains carried *SPC110-mCherry stu2-AID TOR1-1 fpr1Δ* alleles and expressed an ectopic copy of *stu2^ΔBL^ ^ΔCTS^-NLS^SV40^-FRB* either *with NUF2* (M4354) *or NUF2-FKBP* (M4318), to enable tethering of the ectopic FRB-tagged Stu2 to Nuf2 upon rapamycin addition (in the case of *NUF2-FKBP*) or to prevent tethering in the absence of FKBP (in the case of *NUF2* alone). Exponentially growing cells were treated with auxin for 2 h to degrade the endogenous Stu2-AID protein and exhibit a pre-anaphase arrest. After 2 h rapamycin was added (T0) to induce FRB/FKBP dimerization and samples were collected and fixed over a time course of 75 min. Spc110-mCherry and DAPI signals were imaged. The percentage of large-budded cells progressing through anaphase was quantified for each time point, excluding non-large-budded cells from analysis. Bottom left: Representative images of arrested cells and cells that progressed through anaphase. Bottom right: Percentage of the cells that progressed into anaphase after the addition of rapamycin are quantified, based on two biological replicates (n=100-145 per time point for each replicate). Mean and standard deviation are displayed for each time point.

Given this finding, we asked whether restoring Stu2’s localization to different subcellular locations by artificially tethering Stu2^ΔBL^ ^ΔCTS^-NLS^SV40^ to several of Stu2’s interacting partners would restore viability. We utilized an FRB/FKBP rapamycin induced dimerization system (Haruki et al., 2008), tagging Stu2 at the carboxy-terminus with FRB and tagging several of its binding partners with FKBP. Artificially tethering Stu2^ΔBL^ ^ΔCTS^-NLS^SV40^-FRB to well-established Stu2 interactors, including the microtubule plus-end-binding MAPs Bik1 and Bim1 (Stangier et al., 2018; Wolyniak et al., 2006), or the microtubule organizing center component γ-tubulin, Tub4 (Greenlee et al., 2022), did not rescue cell viability. Remarkably, cell viability was restored when this lethal mutant was artificially tethered near its native binding site at the C-terminus of Nuf2 on the kinetochore (Zahm et al., 2021; Fig. 4B and S4B-C).

To further understand why Stu2’s kinetochore function is vital for cell viability, we examined mitotic progression defects in *stu2^ΔBL^ ^ΔCTS^-NLS^SV40^* mutants. Consistent with total loss-of-function mutants (Severin et al., 2001), we found that *stu2^ΔBL^ ^ΔCTS^-NLS^SV40^* mutants arrested in metaphase in a spindle assembly checkpoint (SAC)-dependent manner, suggesting biorientation defects (Fig. S4D-E). Preventing the SAC arrest by deletion of a SAC component, *MAD3*, did not rescue cell viability, indicating that SAC overactivation is not the cause of cellular death (Fig. S4F). Furthermore, the rescue of cell viability observed when tethering Stu2^ΔBL^ ^ΔCTS^-NLS^SV40^-FRB to Nuf2 was not compromised in a *mad3Δ* mutant (Fig. S4G). These data suggest that Stu2 carries out a kinetochore function responsible for proper chromosome segregation. In the absence of this function, kinetochore-microtubule attachments are dysregulated, leading to SAC activation and cell death even if the SAC is bypassed.

To demonstrate that the mitotic arrest observed in *stu2^ΔBL^ ^ΔCTS^-NLS^SV40^* mutants is due to a lack of kinetochore activity, we tested if cells could bypass the arrest by re-tethering Stu2 to Nuf2. Indeed, we found that re-tethering Stu2^ΔBL^ ^ΔCTS^-NLS^SV40^-FRB to Nuf2-FKBP lead to rapid anaphase onset (Fig. 4C). Furthermore, both the rescue of mitotic arrest and cell viability depend on Stu2’s ability to bind tubulin dimers, as a *stu2* mutant with additional point mutations in each of its TOG domains, which disrupt TOG-tubulin interactions (i.e. Stu2^R200A^ ^R519A^ ^ΔBL^ ^ΔCTS^-NLS^SV40^- FRB; Ayaz et al., 2012, 2014), was incapable of rescuing either (Fig. S4H-I).

### Stu2’s precise kinetochore position is critical for function, suggesting a spatial separation mode of action

To understand the role of kinetochore-associated Stu2, we considered the mitotic defects described above and our prior observations that Stu2 confers tension sensitivity to kinetochores in vitro, both stabilizing high-tension attachments and destabilizing low-tension attachments(Miller et al., 2016). Given these observations, it is plausible that Stu2 plays a critical role in the correction of erroneous attachments. Our previous model proposed that Stu2 provides additional microtubule-binding capacity at high tension. However, at low tension, it functions to destabilize attachments, possibly by directly competing with or occluding the microtubule-binding activities of other kinetochore components. This destabilization would be relieved by tension-dependent stretching of the kinetochore structure, spatially separating Stu2 from the other microtubule-binding kinetochore elements (Miller et al., 2016; Fig. 5A).

**Figure 5.**
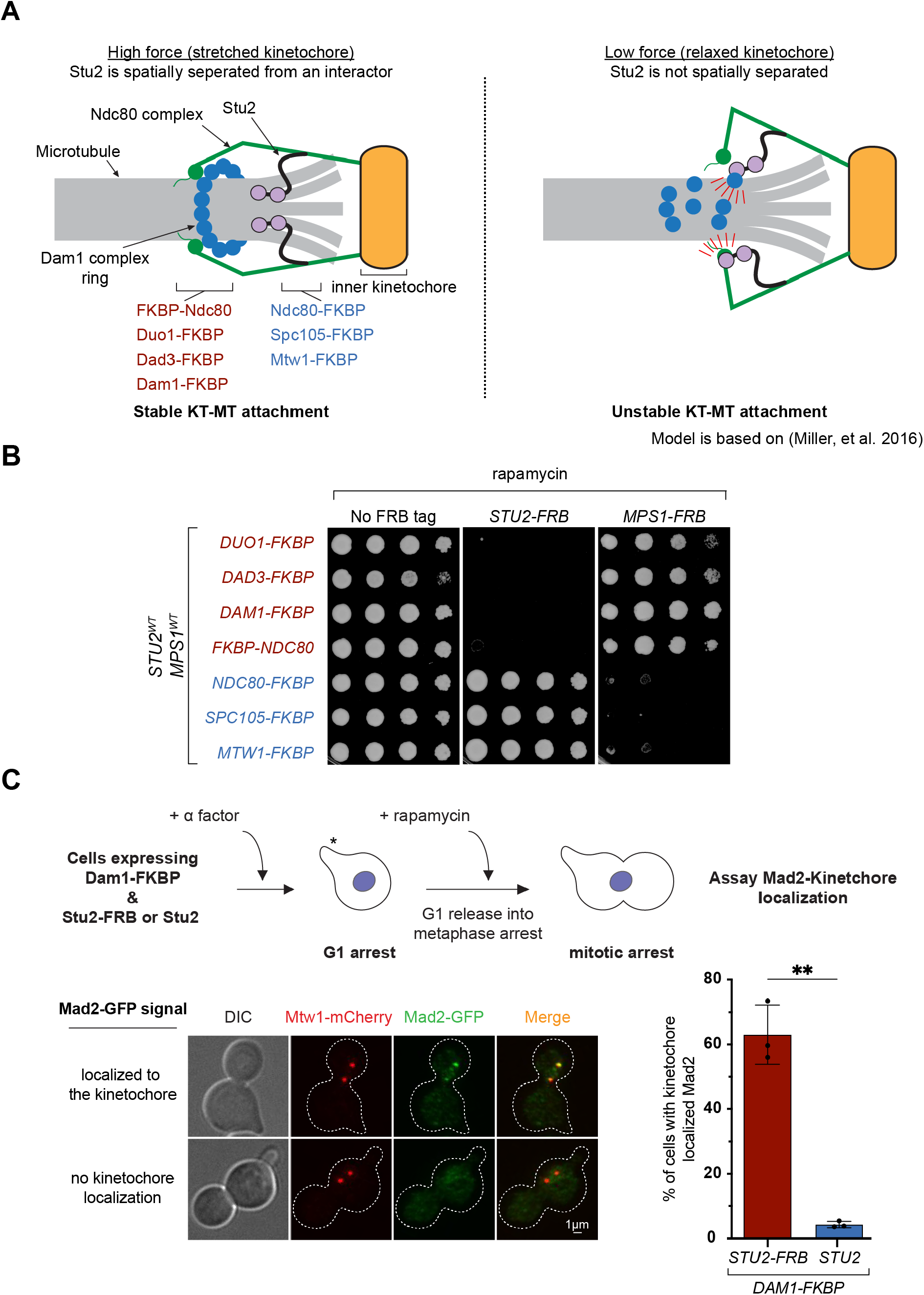
Stu2’s precise kinetochore position is critical for function. (A) A model of how Stu2 might selectively destabilize kinetochore-microtubule attachments at low tension based on (Miller et al., 2016). At high tension, the kinetochore is stretched and Stu2 is spatially separated from a site of interaction. At low tension, the kinetochore is relaxed, and Stu2 can interact with a binding partner, disrupting the kinetochore-microtubule attachment. (B) Cell viability was analyzed by plating five-fold serial dilutions in the presence rapamycin to induce the dimerization of FRB and FKBP. Strains contained endogenous *STU2^WT^* and *MPS1^WT^* with *TOR1-1 fpr1Δ* and an FKBP tagged protein (*DUO1-FKBP*, M5519; *DAD3-FKBP*, M5516; *DAM1-FKBP*, M4784; *FKBP-NDC80*, M5510; *NDC80-FKBP*, M1427; *SPC105-FKBP*, M4613; or *MTW1-FKBP*, M4614) in the absence of an FRB tagged protein (Left), or also containing an ectopic *STU2-FRB* (Middle; with an FKBP tagged protein: *DUO1-FKBP*, M5673; *DAD3-FKBP*, M5643; *DAM1-FKBP*, M5655; *FKBP-NDC80*, M5637; *NDC80-FKBP*, M5631; *SPC105-FKBP*, M5634; or *MTW1-FKBP*, M5646), or with ectopic *MPS1-FRB* (Right; with an FKBP tagged protein: *DUO1-FKBP*, M5675; *DAD3-FKBP*, M5645; *DAM1-FKBP*, M5657; *FKBP-NDC80*, M5639; *NDC80-FKBP*, M5633; *SPC105-FKBP*, M5636; or *MTW1-FKBP*, M5648). DMSO control plates are in Fig. S5. (C) Top: A schematic of the experimental design. Exponentially growing *DAM1-FBKP MAD2- 3GFP MTW1-mCherry pMET-cdc20 TOR1-1 fpr1Δ* cells expressing an ectopic copy of *STU2^WT^* (M6423) or *STU2^WT^-FRB* (M6134) were treated with α factor (mating pheromone) for 3 h to synchronize cells in G1 in synthetic media lacking L-methionine. The asterisk denotes a shmoo. After 3 h, cells were released from G1 arrest into a metaphase arrest by depleting Cdc20 (i.e. in the presence of L-methionine) and in the presence of rapamycin. After 2 h, samples were collected and fixed. Mad2-3GFP and Mtw1-mCherry signals were imaged. Percentage of large- budded cells with a signal of Mad2-3GFP colocalized to Mtw1-mCherry was quantified. Bottom- left: Representative images. Bottom-right: Percentage of cells with Mad2-3GFP colocalized to Mtw1-mCherry are quantified. The graphs represent three biological replicates, n=50-64 per genotype in each replicate. Mean and standard deviation are shown. P value was determined using a two-tailed unpaired *t* test (** = p<0.01).

If this model is correct, the precise positioning of Stu2 would be important for it to carry out its tension-sensitive function. The artificial tethering tools we developed provide a great means to test this hypothesis. We hypothesized that tethering Stu2-FRB at or near its native binding site (the tetramerization junction of the Ndc80 complex (Zahm et al., 2021), specifically the C-terminus of Ndc80 or Nuf2) would be tolerated by the cell. However, moving Stu2 "further out" (i.e. distal to the centromere) by altering the FKBP location would destabilize attachments inappropriately causing cell death.

We tested this by tethering an ectopic copy of full-length Stu2-FRB in the presence of endogenous Stu2. As expected, tethering Stu2 to the C-terminus of Ndc80 was well tolerated. Surprisingly, tethering Stu2 to the N-terminus of Ndc80 resulted in dominant lethality (Fig 5B). We further tested tethering Stu2-FRB to other kinetochore components, both near its native binding site (i.e. the C-termini of Spc105 and Mtw1) and to more centromere-distal parts of the outer kinetochore (C-termini of Dam1, Duo1, and Dad3), observing the same trend (Fig 5B and S5A). As a control, we replicated previous findings by Aravamudhan et al., where Mps1-FRB localization was similarly manipulated. This prior work showed that tethering Mps1 near its substrate, Spc105, triggered a hyperactive SAC and cell death (Aravamudhan et al., 2015; Parnell et al., 2024), while centromere-distal tethering had no impact on cell viability. Notably, Stu2 and Mps1 exhibited completely opposite phenotypes when ectopically tethered (Fig 5B and S5A).

If our spatial separation model is correct, the lethality caused by tethering Stu2-FRB to the N-terminus of Ndc80 and C-termini of Dam1, Duo1, and Dad3 should result from the inability to form stable kinetochore-microtubule attachments. To test this, we monitored unattached kinetochores by assessing recruitment of the SAC components Mad2 or Bub1 in response to Stu2 tethering to the C-terminus of Dam1. Cells expressing Mad2-GFP Mtw1-mCherry and either wild-type *STU2* or *STU2*-*FRB* were released from a G1 arrest into rapamycin-containing media and subsequently arrested in metaphase by depleting Cdc20. Consistent with the hypothesis above, tethering Stu2- FRB to Dam1 resulted in a substantial number of cells exhibiting Mad2 kinetochore recruitment (Stu2-FRB: 63.0 ± 9.2%; Stu2: 4.2 ± 1.0 %; Fig. 5C). Similarly, tethering Stu2-FRB to Dam1 resulted in the recruitment of Bub1-GFP to the kinetochore (Fig. S5B). This suggests that mis- localizing Stu2 closer to the other microtubule-binding elements of the outer kinetochore overrides potential tension-sensitive mechanisms, leading to inappropriate destabilization of kinetochore-microtubule attachments.

## DISCUSSION

Previous work demonstrated that the ch-TOG family of proteins function as critical regulators of microtubules by controlling polymerization, depolymerization, nucleation, and kinetochore-microtubule interactions (Brouhard et al., 2008; Chen et al., 1998; Flor-Parra et al., 2018; Fox et al., 2014; Geyer et al., 2018; King et al., 2020; Kinoshita et al., 2002; Kosco et al., 2001; Miller et al., 2016; Ohkura et al., 2001; Podolski et al., 2014; Thawani et al., 2018; Usui et al., 2003; Widlund et al., 2011). Here, we make a number of important mechanistic insights into Stu2’s essential function, which can be summarized in three main points: 1) by developing a separation of function mutant, we show that Stu2’s ability to regulate microtubule dynamics is dispensable for cell viability; 2) using an inducible artificial tethering rescue system, we demonstrate that Stu2 has a kinetochore function essential for proper chromosome segregation and cell viability; 3) the precise kinetochore localization of Stu2 is critical – ectopically tethering Stu2 to the most centromere-distal kinetochore components leads to the accumulation of unattached kinetochores and cell death. Based on our findings, we speculate that Stu2’s ability to confer tension sensitivity to kinetochores is essential for establishment of bioriented kinetochore attachments in cells. Previous work has also suggested that other members of this larger protein family, such as the human ortholog ch-TOG and the fission yeast Dis1 and Alp14, play important roles in kinetochore function (Herman et al., 2020; Yukawa et al., 2019). However, due to the pleiotropic defects of loss-of-function mutants, it has been challenging to understand the importance of their kinetochore roles. Our findings highlight the significance of the budding yeast, Stu2’s, kinetochore role and provide insights into the molecular mechanisms by which it performs its various activities. This has significant implication on our understanding of the role of ch-TOG proteins in chromosome segregation.

### Stu2 carries an essential kinetochore function independent of its ability to polymerize microtubules

Stu2 acts as both a microtubule polymerase and depolymerase, which relies on its array of TOG domains and basic linker domain. The TOG domains bind to tubulin dimers, facilitating their addition to a growing microtubule tip or their removal from a shrinking tip, while the basic linker enables Stu2 to associate with the negatively charged microtubule lattice. Previous studies have shown that at least one active TOG domain is necessary for cell viability, as is a conserved 15- residue patch within the basic linker (Ayaz et al., 2014, 2012; Geyer et al., 2018; Herman et al., 2020; Miller et al., 2019). Here, we demonstrate that this conserved patch also contains Stu2’s nuclear localization sequence (Fig. 2). Surprisingly, a complete deletion of the basic linker can sustain viability when fused to an exogenous NLS (*stu2^ΔBL^ -NLS^SV40^*; Fig. 3A). However, Stu2^ΔBL^ - NLS^SV40^ is deficient in regulating microtubule dynamics (Fig. 3B-D). Finally, we identified a *stu2* mutant (*stu2^ΔBL^ ^ΔCTS^ -NLS^SV40^*) that supports cell viability when artificially tethered near its native kinetochore binding site but is otherwise lethal (Fig. 4B). Together, these data indicate that Stu2 possesses a kinetochore function essential for cell viability, likely independent of its ability to polymerize microtubules (also see Supplementary Discussion Point 2).

### Insights into Stu2’s kinetochore function

Similar to null mutants of Stu2 (Severin et al., 2001), *stu2^ΔBL^ ^ΔCTS^ -NLS^SV40^* mutants arrest prior to anaphase in a SAC-dependent manner. Cells overcome this arrest, presumably by satisfying the SAC, when Stu2^ΔBL^ ^ΔCTS^ -NLS^SV40^ is tethered to the kinetochore to fulfill an essential function (Fig. 4C). We are aware of three potential kinetochore functions of Stu2 which, in their absence, could result in chromosome biorientation defects, SAC activation, and viability defects. The first is kinetochore capture. In this capacity, Stu2 aids in the capture of kinetochores by nucleating kinetochore derived microtubules and regulating microtubule dynamics during lateral attachment. However, evidence suggests that both functions are ultimately dispensable for kinetochore capture and cell viability (Gandhi et al., 2011; Kitamura et al., 2010; Vasileva et al., 2017). Second, kinetochore-associated Stu2 may regulate the dynamics of end-on attached microtubules. The idea that this function is essential for cell viability, however, is inconsistent with a number of our observations. Two different *stu2* mutants defective in their ability to stimulate microtubule polymerization and depolymerization, Stu2^ΔBL^-NLS^SV40^ identified in this study and Stu2^ΔTOG2^ (Geyer et al., 2018), can still sustain cell viability. Furthermore, our prior work showed no difference in the dynamics of kinetochore-bound microtubules in the presence or absence of Stu2 in vitro (Miller et al., 2016), implying that kinetochore-associated Stu2 may not contribute to regulating microtubule dynamics when bound to end-on attached kinetochores. The third function is Stu2’s role in conferring tension sensitivity to kinetochores. In previous work using purified kinetochores in vitro, we found that Stu2 imparts this tension sensitivity through dichotomous activities. At high tension, Stu2 stabilizes attachments, while at low tension, its presence leads to attachment destabilization (Miller et al., 2016). We proposed a spatial separation model to explain these functions. At high tension, Stu2 acts as a microtubule-binding element that ensures kinetochore-microtubule attachments are robust. Conversely, at low tension, relaxation of kinetochore stretching enables Stu2 to compete with other kinetochore-microtubule binding components, thereby disrupting kinetochore-microtubule attachments. This attachment destabilization may occur through interference with Ndc80’s CH domain microtubule binding, inhibiting Dam1 complex oligomerization or microtubule binding, or via other mechanisms.

Here, we tested this spatial separation model by ectopically tethering Stu2 to the centromere-distal, N-terminal tail of Ndc80 and C-termini of Dam1 complex components (Duo1, Dad3, and Dam1) and showed that this led to cell death and recruitment of the SAC components Mad2 and Bub1, indicating unattached kinetochores (Fig. 5 and Fig. S5). While further work is needed to dissect the mechanistic basis of Stu2’s kinetochore functions, this work demonstrates that the precise positioning of Stu2 within the kinetochore is crucial, supporting the spatial separation model described above. It is also worth noting that other modes of action might explain our data. For instance, Stu2’s TOG domains may sense the conformational state of the microtubule plus tip, selectively binding to certain tip conformations. When the TOG domains are not bound to the microtubule tip, they may be free to interact with other factors that disrupt the kinetochore-microtubule attachment. This idea aligns with our previous in vitro observations using purified kinetochores, where Stu2’s activity conferred microtubule tip state dependence: Stu2 specifically disrupted kinetochore-microtubule attachments at low tension on disassembling, but not on assembling, microtubules (Miller et al., 2016). If that model is correct, perhaps tethering Stu2 to the N-terminal tail of Ndc80 (or to various centromere-distal C-termini of the Dam1 complex) “sequesters” it away from the microtubule tip, thus freeing the TOG domains to disrupt the kinetochore-microtubule attachment. Ultimately, it is possible that a combination of spatial separation and microtubule tip conformation sensing is at play.

## ACKNOWLEDGEMENTS

The authors declare no competing financial interests. We thank Sue Biggins, Trisha Davis, Eris Duro and Adèle Marston for reagents; Sue Biggins, Jake Herman, Trisha Davis and the Miller lab for critical reading of the manuscript. This work was supported by funds from the Judy Chandler Webb Endowed Chair in the Biological Sciences (to G.B), Robert A Welch Foundation I-1901 (to L.R.), 5 For the Fight (to M.P.M), Pew Biomedical Scholars (to M.P.M), and NIH grants R01GM098542 (to L.R.), R01GM47842 (to G.B.), and R35GM142749 (to M.P.M).

## METHODS

Yeast strains and plasmids *Saccharomyces cerevisiae* strains used in this study are described in Table S1 and are derivatives of M3 (W303) or M1652 (S288C). Standard media and microbial techniques were used and yeast strains were constructed by standard genetic techniques (Rothstein, 1991; Sherman, 1991). *NUP2-mKATE2*, *KAP60-3FLAG, BIK1-2xFKBP12*, *BIM1-2xFKBP12*, *TUB4- 2xFKBP12*, *DUO1-2xFKBP12*, *DAD3-2xFKBP12*, *DAM1-2xFKBP12*, *2xFKBP12-NDC80*, *NDC80-2xFKBP12*, *SPC105-2xFKBP12*, and *MTW1-2xFKBP12* were constructed by PCR-based methods (Longtine et al., 1998). Strains containing previously described alleles were also generously provided (*cdc20-IAA17* by Eris Duro and Adèle Marston, *mad3Δ*, *MTW1-3GFP*, *MTW1-mCherry*, *pMET-cdc20*, *MAD2-3GFP*, and *BUB1-GFP* from Sue Biggins, *GFP-TUB1* and *cdc23-1* from Georjana Barnes and David Drubin, and *SPC110-mCherry* from Trisha Davis). *stu2- 3HA-IAA7*, *pGPD1-OsTIR1* and *pSTU2-STU2-3V5* are described in (Miller et al., 2016) and *NUF2-2xFKBP12* described in (Zahm et al., 2021). *STU2-3V5* variants were constructed by mutagenizing pM225 as described in (Liu and Naismith, 2008; Tseng et al., 2008). *STU2-Halo- 3V5* variants were constructed by mutagenizing pM630 as described in (Liu and Naismith, 2008; Tseng et al., 2008). *pSTU2-STU2-GFP* is described in (Miller et al., 2019). *STU2-GFP* variants were constructed by mutagenizing pM488 as described in (Liu and Naismith, 2008; Tseng et al., 2008). *stu2^592-607^-GFP-GST* variants were constructed by mutagenizing pM774 as described in (Liu and Naismith, 2008; Tseng et al., 2008). *pSTU2-STU2-FRB-3V5* is described in (Zahm et al., 2021). *STU2-FRB-3V5* variants were constructed by mutagenizing pM531 as described in (Liu and Naismith, 2008; Tseng et al., 2008). *pMPS1-MPS1-FRB-3V5* was constructed by cloning *pMPS1-MPS1* into an FRB-3V5 plasmid as described in (Liu and Naismith, 2008; Tseng et al., 2008). Plasmids and primers used in construction are listed in Table S2 and further details of plasmid construction including plasmid maps are available upon request.

### Auxin inducible degradation

The auxin inducible degron (AID) system was used as described in (Nishimura et al., 2009). Cells expressed C-terminal fusions of the protein of interest to an auxin responsive protein (IAA7) at the endogenous locus. Cells also expressed *TIR1*, which is required for auxin-induced degradation. 500 μM auxin (Sigma-Aldrich, Cat#I3750-5G-A) dissolved in DMSO was added to liquid media or top-plated on agar to induce degradation of the AID-tagged protein. Auxin was added to liquid media 2 h prior to harvesting cells when arresting cells in mitosis or otherwise 30 min prior to harvesting.

### FRB/FKBP tethering system

For the cell viability assay, rapamycin (to induce FRB/FKBP dimerization; LC Laboratories, Cat#R5000) and auxin (to degrade Stu2-AID) dissolved in DMSO were top-plated on agar plates to a final concentration of 500 ng/ml rapamycin and 500 µM auxin. For the liquid culture experiments a final concentration of 500 ng/ml rapamycin was used with or without the addition of 500 µM of auxin (as indicated in the figure legend for each experiment) for the indicated time periods.

α factor arrest and release into *pMET-cdc20* mitotic arrest Cells carrying *pMET-cdc20* were maintained and grown on synthetic media lacking L- Methionine. α factor (University of Utah Core Synthesis Facility) was added at a concentration of 1 µg/ml for 3 h to arrest cells in G1. To release cells, the culture was pelleted and washed 3 times (resuspending and pelleting again each time) with YPAD media + 1% DMSO. After the third wash, cells were resuspended in YPAD media + 500 ng/ml rapamycin + 8 mM L- Methionine to repress *pMET-cdc20* expression and arrest cells in metaphase. 8 mM of L- Methionine was subsequently added each hour.

### Spot viability assay

For the spot viability assay, the desired strains were grown overnight on YPD plates containing yeast extract peptone plus 2% glucose (YPD) medium. The following day, equal amounts of each desired strain were used for a 1:5 dilution series and were spotted on YPD + DMSO, YPD + 500 μM auxin, YPD + DMSO + 6.5µg/mL benomyl, YPD + 500 μM auxin + 6.5µg/mL benomyl, YPD 500 ng/ml rapamycin, or YPD + 500 μM auxin + 500 ng/ml rapamycin plates. Plates were incubated at 23°C for 2–3 days unless otherwise specified.

### Cell Fixation, Imaging conditions, and Image Analysis

Exponentially growing cultures were treated with 500 μM auxin for 2 h prior to harvesting when arresting cells in mitosis or 30 min prior to harvesting when arresting cells in G1. For time course experiments, exponentially growing cells were treated for 2 h with 500 μM auxin treatment, followed by addition of 500 ng/ml rapamycin (timepoint 0 min), and samples were harvested over time as indicated for each experiment. Harvested samples were then fixed (see below) and analyzed for Stu2-GFP localization and Nup2-mKATE localization, Mtw1-GFP distribution and spindle length and orientation (Spc110-mCherry), or determined to be mononucleate or binucleate. For each strain, an aliquot of cells was fixed with 3.7% formaldehyde in 100mM phosphate buffer (pH 6.4) for 1-3 min (consistent within every experiment). Cells were washed once with 100 mM phosphate (pH 6.4), resuspended in 100 mM phosphate, 1.2 M sorbitol buffer (pH 7.5) and permeabilized with 1% Triton X-100 stained with 1 μg/ml DAPI (4’, 6-diamidino-2- phenylindole; Molecular Probes). Cells were imaged using a DeltaVision Ultra microscope with a 60X objective (NA = 1.42), equipped with a sCMOS digital camera. Either twenty-one or fifteen (consistent within every experiment) Z-stacks (0.2 micron apart) were acquired, and all frames were deconvolved using standard settings. For quantification of nuclear localization, image stacks were projected using the average intensity. For observation of Mtw1-GFP or Spc110-mCherry, image stacks were maximally projected. softWoRx image processing software was used for image acquisition and processing.

A software was developed to automate the analysis of cellular GFP localization as well as spindle lengths. A cell segmentation model was used to segment individual cells within DIC images. For this implementation, the model developed by [https://github.com/alexxijielu/yeast_segmentation], which uses Mask-RCNN Segmentation [https://arxiv.org/pdf/1703.06870.pdf], was used; however, due to the modular nature of the software, any segmentation model could be easily used. After the objects (i.e. individual mother and daughter cells) are individually segmented, mother-daughter pairs are merged together by calculating the nearest neighbor of each object within 3 pixels. If the nearest neighbor of object A is object B, and the nearest neighbor of object B is object A, then the two objects are merged as a single mother-daughter pair, representing a large-budded cell; otherwise, the cells are ignored during further analysis. Segmented cells that were improperly segmented were manually removed from analysis. Once the large-budded cells are detected in the DIC image, the outlines of those large-budded cells are used to extract the location of each cell in the mCherry/mKATE and GFP images.

The spindle length is calculated by first identifying the locations of high intensity Spc110- mCherry signal for a given large-budded cell. This is accomplished by converting the mCherry image for a given cell to grayscale, smoothing the image with a Gaussian blur [kernel size of (3,3) and kernel standard deviation of 1 along both the x and y axes] and then cleaning the noise from the image by using an adaptive Gaussian threshold with Otsu binarization. Next, contours in the image are detected and the two largest contours are extracted. To determine the spindle length, the moment of each contour is calculated, and then the center of each contour is calculated from the moment. This center point represents the spindle pole body in the Spc110-mCherry images. The spindle line is then calculated as the line between the center of each contour. Length of this spindle line is calculated and represents the spindle length.

To calculate the nuclear and cytoplasmic GFP intensities, the contours of the high intensity Nup2-mKATE signal, representing the nucleus, is determined as described above (similar to determining the contours of the Spc110-mCherry signal). Next, the background data is removed from the GFP image by using a rolling ball background subtraction algorithm with a radius of 50. The nuclear GFP intensity is then determined by adding the intensity values from each point in the GFP image that falls within the nucleus contour determined earlier in this process. The final step is to calculate the cellular intensity. Similar to the previous process, the GFP image is converted to grayscale, smoothed with a Gaussian blur [kernel size of (13, 13) and kernel standard deviation of 5 in both the x and y axes], and cleaned using an adaptive Gaussian threshold with Otsu binarization. Contours are then extracted from the image. The cellular GFP intensity is then calculated by extracting the intensity values of every point within the cellular contour. Cytoplasmic GFP intensity is calculated by subtracting the nuclear GFP intensity from the cellular GFP intensity.

To analyze spindle orientation and Mtw1-GFP distribution, projected images were imported into FIJI - ImageJ (NIH) for manual analysis. To determine spindle orientation, the angle tool was used to measure the angle between the spindle axis (a line between the two Spc110- mCherry puncta) and the mother-daughter bud axis (a line between the center of the distal ends of the large-budded cell). Values were reported as less than or greater than 45 degrees. To determine if a cell had a bi-lobed versus mono-lobed kinetochore cluster, Mtw1-GFP signals were examined. A mono-lobed kinetochore cluster was identified as an Mtw1-GFP puncta that could not be distinguished into two separate punctae, whereas bi-lobed kinetochore clusters had two distinguishable Mtw1-GFP punctae. Cells belonging to each group were counted and represented as a proportion for each strain. To determine if a cell had a detached kinetochore, Mtw1-GFP signals were examined. A detached kinetochore cluster was identified as an Mtw1-GFP puncta that was dimmer and distinct from the intra-spindle Mtw1-GFP signal. Cells belonging to this group versus cells with no detached kinetochores were counted and represented as a proportion for each strain.

To determine the proportion of mononucleate vs binucleate cells within the large budded cell population, images were imported into FIJI for manual analysis. Cells with a bud roughly at least half the diameter of the mother cell were considered large-budded. DAPI and Spc110- mCherry signals were used to determine if a cell was binucleate or mononucleate.

To analyze Mad2-GFP signal at the kinetochore, images were imported into FIJI for manual analysis. Cells with a bud roughly at least half the diameter of a mother cell that had a shmoo were analyzed. Cells with Mtw1-mCherry signals further apart more than 2 µm were neglected. Mtw1-mCherry signal was circled and then the Mad2-GFP signal in the highlighted area was quantified. Due to Mad2-3GFP background, a cut off value of maximum Mad2-3GFP signal was used to differentiate positive vs negative signals. Cells were considered positive (Mad2 localized to the kinetochore) if Mad2 signal had a maximum higher than 15,000 AU and was considered negative if the maximum value was less than 15,000 AU. If a cell had an Mtw1- mCherry signal detached from the spindle (unattached kinetochore) with a colocalized Mad2-GFP signal with a maximum stronger than 15,000 AU it was counted as positive.

To analyze Bub1-GFP signal at the kinetochore, images were imported into FIJI for manual analysis. Cells with a bud roughly at least half the diameter of a mother cell that had a shmoo were analyzed. Cells with Mtw1-mCherry signals further apart more than 2 µm were neglected. Cell with no Bub1-GFP signals colocalized to Mtw1-mCherry were assigned as no Bub1 signal; cells with one or two Bub1-GFP foci, both of which are weaker than a cut off value of 80,000 AU (Bub1-GFP signal was circled in FIJI and measured), were assigned as weak signal; while those with at least one puncta of Bub1-GFP that was stronger than 80,000 AU were assigned to have a strong Bub1 signal.

### Live cell imaging conditions and analysis

Exponentially growing cultures, grown in synthetic complete media, were treated with 500 μM auxin 30 min prior to harvesting to degrade Stu2-AID and analyzed for Stu2-GFP (ectopically expressed, *STU2^WT^-GFP* or *stu2^ΔBL^-GFP*) distribution and spindle length. For each strain, an aliquot of cells was pelleted and resuspended in a volume of synthetic complete media with 500 μM auxin to optimize cell density for imaging (OD_600_≈5). Cells were adhered to a coverslip coated in Concanavalin A as described in (Fees et al., 2017) and the chamber was sealed using petroleum jelly. Cells were imaged using a DeltaVision Ultra microscope with a 60X objective (NA = 1.42), equipped with a sCMOS digital camera. Images of the Spc110-mCherry signal, Stu2-GFP signal and DIC were acquired through the thickness of the cells using Z-stacks 0.3 micron apart. These images were acquired every 1 min. All frames were deconvolved using standard settings. Image stacks were maximally projected for analysis of spindle lengths and Stu2 distribution. softWoRx image processing software was used for image acquisition and processing. Projected images were imported into FIJI for analysis. Background signal was subtracted, and in cells undergoing spindle elongation, the distance between Spc110-mCherry punctae was measured every min beginning 5 min prior to anaphase onset until 20 min post-anaphase onset. Spindle length at anaphase onset was determined for a given cell by selecting the spindle length at the time point prior to which an increase in spindle length of at least 0.2 microns was observed and followed by an increase in spindle length over the next 3 time points. Maximum rate of spindle elongation for a given cell was calculated by determining the maximum difference in spindle length over a 2-min time period and calculating the rate of spindle elongation over that time period.

### Immunoprecipitationsh

An α-Flag immunoprecipitation of Kap60-3Flag was performed (essentially as described in (Akiyoshi et al., 2010) to purify Kap60 and associated binding partners from asynchronously growing *S*. *cerevisiae* cells as described below. Strains expressing *stu2-AID* were grown overnight in yeast peptone dextrose rich (YPD) medium and back diluted to OD_600_≈0.4 in the morning. 30 min prior to harvest, once cells reached OD_600_≈2-3, 500 μM auxin was added to the culture to degrade Stu2-AID. Protein lysates were prepared by lysing cells in a Freezer Mill submerged in liquid nitrogen. Lysed cells were resuspended in buffer H (BH; 25 mM HEPES pH 8.0, 2 mM MgCl_2_, 0.1 mM EDTA, 0.5 mM EGTA, 0.1% NP-40, 15% glycerol with 150 mM KCl) containing protease inhibitors (at 20 μg/mL final concentration each for leupeptin, pepstatin A, chymostatin and 200 μM phenylmethylsulfonyl fluoride) and phosphatase inhibitors (0.1 mM Na-orthovanadate, 0.2 μM microcystin, 2 mM β-glycerophosphate, 1 mM Na pyrophosphate, 5 mM NaF) followed by ultracentrifugation at 98,500 xg for 90 min at 4°C. Dynabeads conjugated with α-Flag were incubated with extract for 3 h with constant rotation, followed by three washes with BH containing protease inhibitors, phosphatase inhibitors, 2 mM dithiothreitol (DTT) and 150 mM KCl. Beads were further washed twice with BH containing 150 mM KCl and protease inhibitors. Associated proteins were eluted from the beads by incubation in 1x SDS sample buffer at 95°C for 3 min.

### Immunoblotting

For immunoblot analysis, cell lysates were prepared as described above. Standard procedures for sodium dodecyl sulfate-polyacrylamide gel electrophoresis (SDS-PAGE) and immunoblotting were followed as described in (Burnette, 1981; Towbin et al., 1979). A nitrocellulose membrane (Bio-Rad) was used to transfer proteins from polyacrylamide gels. Commercial antibodies used for immunoblotting were as follows: Mouse monoclonal α-FLAG (Sigma-Aldrich, Cat#F3165) 1:3,000; Mouse monoclonal α-V5 (Invitrogen, Cat#R960-25) 1:5,000; Mouse monoclonal α-Pgk1 (Abcam, Cat#22C5D8) 1:5,000; Mouse monoclonal α-HA (Roche, 12CA5; Cat#11583816001) 1:2500; Rabbit Polyclonal α-FKBP12 (Invitrogen, Cat#PA1-026A) 1:1000. The secondary antibodies used were sheep α-mouse (HRP) (Cytiva/GE Biosciences, Cat#NA931-1ML) and Donkey α-rabbit (HRP) (Cytiva/GE Biosciences, Cat#NA934-1ML) at a 1:10,000 dilution. Antibodies were detected using the SuperSignal West Dura Chemiluminescent Substrate (Thermo Scientific) and ProSignal Femto substrate (Genesee Scientific).

### Statistics

GraphPad Prism version 9.1 was used for statistical analysis. Data normality was assumed for all experiments. Student’s *t* test was used for comparisons between groups in experiments with 2 groups, an ordinary one-way ANOVA followed by a Tukey’s multiple comparisons test in experiments with more than two groups, and Fisher’s exact test was used for categorical data. In the text, mean ± SD is reported.

### Multiple sequence alignments

Fungal proteins related to *Saccharomyces cerevisiae* Stu2 were identified using a PSI-BLAST (Altschul et al., 1997) search on NCBI. Multiple sequence alignments of the entire proteins were generated with ClustalOmega default parameters and displayed in JalView 1.8.

### Whole cell lysate polymerase assay

All strains were grown in standard rich media (YPD) at 25°C since they contained a temperature-sensitive allele. Cells were harvested as previously described in (Bergman et al., 2019). In short, strains were grown overnight and then diluted back into two identical 2L cultures of YPD with 2% glucose. At an OD_600_≈0.4, cultures were then shifted to the non-permissive temperature of 29°C for a 3 h arrest. Strains with AID degrons were treated with 250 µM auxin (3-indole acetic acid; Sigma-Aldrich) in DMSO 30 min before harvesting, 2.5 h into the arrest. Cells were then harvested by serial centrifugation at 6000 rpm in a Sorvall RC5B with a SLA-3000 rotor for 10 min at 4°C. Cell pellets were then resuspended in cold ddH_2_O and pelleted in a Hermle Z446K centrifuge for 3 min at 4000 rpm 4°C. This was repeated twice more, with the last spin (omitting the wash) to remove all standing water from the pellet via aspiration. The cells were then snap-frozen in liquid nitrogen and stored at -80°C.

As previously described, approximately 5g of frozen cell pellet was weighed in a pre-chilled medium-sized SPEX freezer mill vial. The pre-chilled chamber was then milled (submerged in liquid nitrogen) following a protocol that consisted of a 3 min pre-chill, then 10 cycles of 3 min grinding at 30 impacts per second (15 cps) and 1 min of rest. The resulting powered lysate was stored at -80°C.

As previously described in (Bergman et al., 2019), previously purified bovine tubulin was cycled to remove nonfunctional tubulin and any remaining impurities. This tubulin was mixed with both biotin-conjugated and rhodamine labeled porcine tubulin (Cytoskeleton Inc.). The tubulin mix was then resuspended in PEM buffer (80 mM PIPES pH 6.9, 1 mM EGTA, 1 mM MgCl2) to a final concentration of 5 mg/mL unlabeled tubulin, 1 mg/mL biotin-labeled tubulin, and 1 mg/mL far-red labeled tubulin. To stabilize the seeds, GMPCPP (Jena Bioscienes) was added to a final concentration of 1 mM. Aliquots were then snap-frozen in liquid nitrogen and stored at -80°C.

Both glass side preparation and passivation of coverslips was similar to the protocol previously described in (Bergman et al., 2019). To prepare the glass slides for assembly into a flow chamber, microscope slides (Corning Inc.) were washed in acetone for 15 min and then 100% ethanol for 15 min. These slides were left to air dry before being stored in an airtight container.

To prepare the coverslips for assembly into a flow chamber, cover glasses (1.5 thickness, Corning Inc.) were first cleaned by sonication in acetone for 30 min. The coverslips were then soaked for 15 min in 100% ethanol. After 3 thorough, but short, rinses in ddH_2_O, the coverslips were submerged for 2 h in 2% Hellmanex III solution (Hellma Analytics). After this, they were again rinsed in ddH_2_O 3 times. Before proceeding to passivation, the coverslips were blown dry with nitrogen gas. For passivation, a solution containing 0.1 mg/mL pLL-g-PEG:PEG(3.4)-biotin (50%:50%) (SuSoS AG) in 10 mM HEPES was prepared. 50 µL drops were then placed on Parafilm in a humid chamber and coverslips were gently placed onto the drops. After 1 h, the passivated coverslips were washed for 2 min in PBS and rinsed in ddH_2_O for 1 min. Again, the coverslips were blown dry with nitrogen gas and stored in an airtight container at 4°C. These passivated coverslips were used for up to a maximum 3 weeks.

Preparation and assembly of the flow chamber for use in the TIRF based dynamics assay was exactly the same as described in (Bergman et al., 2019).

Similar to (Bergman et al., 2019), 0.22 g of powdered lysate was weighed in a 1.5 mL eppendorf tube pre-chilled in liquid nitrogen. To the powder, 12.5 µL of cold 10X PEM (800 mM PIPES pH 6.9, 10 mM MgCl_2_, 10 mM EGTA), 0.5 µL of Protease Inhibitor Cocktail IV (Calbiochem), and 2.5 µL of 10 µM Halo-conjugated JF646 dye in DMSO was added, spun down briefly, and incubated on ice for 10 min. Lysate was added to pre-chilled polycarbonate ultracentrifuge tubes and cleared of insoluble material by spinning at 34,600 xg for 25 min at 4°C. After ultracentrifugation, 32 µL of cleared soluble lysate was flowed into the chamber prepared previously (see above).

After clarified soluble lysate was added to the prepared chamber, the slides were loaded onto a Nikon Ti2-E inverted microscope with an Oko Labs environmental chamber pre-warmed to 28°C. Images were acquired with a Nikon 60X CFI Apo TIRF objective (NA 1.49) and an Orca Fusion Gen III sCMOS camera (Hamamatsu) at 1.5X magnification using the Nikon NIS Elements software. Using a LUNF 4-line laser launch (Nikon) and an iLas2 TIRF/FRAP module (Gataca Systems) total internal reflection fluorescence illuminated a single focal plane of the field and was imaged every 5 seconds for 30 min.

Analysis was done as described previously in (Bergman et al., 2019). Imaging data was analyzed using FIJI. Registration to correct for stage drift was applied to the raw data (Preibisch et al., 2010; Thévenaz et al., 1998). Kymographs were generated from all microtubules whose entire length was trackable for the entire movie. Kymographs were excluded if the microtubules were crossed or were bundled. For analysis, data from independent technical trials and biological replicates from one genotype were pooled, unless otherwise indicated.

### Protein expression and purification

αβ-tubulin from yeast was overexpressed in *S. cerevisiae* and purified using Ni-affinity and ion exchange chromatography, as previously described in (Geyer et al., 2015) but with minor modifications. Stu2^KR/AA^ mutations were introduced into the Stu2^WT^ background using QuikChange (Stratagene) mutagenesis. All Stu2 constructs were expressed in *E. coli* using Arctic Express Cells, and were induced with 0.5 mM IPTG at 10°C for 24 hrs. Cell pellets were resuspended in lysis buffer (50 mM sodium phosphate dibasic and monobasic mix, 300 mM NaCl, 40 mM imidazole, 5% glycerol with PMSF added to 1 mM final concentration), and lysed using a microfluidizer. The lysate was then clarified by centrifugation before loading onto a His60 Superflow Column (Clontech) and eluted in lysis buffer containing 300 mM imidazole. Pooled elution fractions were then loaded onto a Strep-Tactin Superflow column (IBA, Germany) and eluted in RB100 (25 mM Tris pH 7.5, 100 mM NaCl, 1 mM MgCl_2_, 1 mM EGTA) containing 10 mM desthiobiotin. For storage, eluted samples were exchanged into RB100 with 2 mL, 7K MWCO Zeba spin desalting columns (Thermo Scientific).

### In vitro microtubule dynamics assay

Assay chambers were assembled from slides, coverslips, and double sticky tape as previously described (Geyer et al., 2015). Chambers were preincubated with neutravidin (5 µg/ml) and blocked with 1% F-127 Pluronic acid for 10 and 5 min respectively. The chambers were then rinsed with BRB80 (80 mM PIPES pH 6.9, 1 mM MgCl_2_, 1 mM EGTA) followed by 10 min incubation with GMPCPP seeds made from brain tubulin (5% biotinylated, PurSolutions). Chambers were rinsed again with BRB80 and samples containing αβ-tubulin and accessory protein (Stu2^WT^ or Stu2^KR/AA^) in 1× PEM + 0.1 mg/ml bovine serum albumin (BSA) + 1 mM GTP were flowed into the same and immediately sealed with VALAP. Microtubule dynamics were measured by time-lapse DIC microscopy, as described previously in (Geyer et al., 2015).

## ONLINE SUPPLEMENTAL MATERIAL

Supplementary Discussion Point 1: a discussion about why Stu2^ΔCTS^ is viable. Supplementary Discussion Point 2: an extended discussion of the evidence indicating that Stu2’s microtubule polymerization ability is not essential for cell viability. Fig. S1 shows conservation of the entire basic linker region of Stu2 across 9 fungal species as well as viability assays of various mutants within Stu2^592-606^, related to Fig. 1. Fig. S2 shows the ratio of nuclear to cytoplasmic GFP signal intensity in Stu2^592-607^-GFP-GST with a copy of GFP that is unable to dimerize and modulating nuclear levels of Stu2^KR/AA^ affects viability, related to Fig. 2. Fig. S3 shows that basic linker domain is required for Stu2’s ability to stimulate microtubule assembly and disassembly in vitro but is dispensable for cell viability as long as Stu2 is localized to the nucleus, related to Fig. 3. Fig. S4 shows that Stu2’s kinetochore function is required for cell viability independently of the SAC and tethering a Stu2 lethal mutant specifically to Nuf2 but not to other interactors rescues cell viability, related to Fig. 4. Fig. S5 shows that tethering Stu2-FRB to Ndc80’s N-terminal tail or to Dam1 components causes cell lethality and Bub1-GFP kinetochore localization when Stu2 is tethered to Dam1, related to Fig. 5.

## SUPPLEMENTARY DISCUSSION

<u>Supplementary discussion point 1</u>: Here, we show that the combined loss of Stu2’s basic linker region and the CTS (Stu2^ΔBL^ ^ΔCTS^ -NLS^SV40^) is lethal unless artificially tethered to the kinetochore. Interestingly, we previously found that although *stu2^ΔCTS^* mutants exhibit dramatically reduced kinetochore co-immunoprecipitation, they show only partial viability defects rather than lethality (Miller et al., 2019; Zahm et al., 2021). If Stu2’s kinetochore localization is essential, why is this the case? One possible explanation is that Stu2 may interact with the kinetochore through an additional element beyond its CTS. This interaction might allow enough Stu2 to localize at the kinetochore to fulfill its essential role, accounting for the moderate cell viability observed in the *stu2^ΔCTS^* mutant. However, this interaction may be too weak to withstand the stringent conditions of biochemical purification in our kinetochore IPs. There are two possible explanations for why the combined loss of the basic linker and CTS is lethal (i.e. *stu2^ΔBL^ ^ΔCTS^ -NLS^SV40^* mutant). The first is that the basic linker itself contains an additional kinetochore-binding element for Stu2. The second possibility is that deletion of the basic linker creates a hypomorphic mutant that cannot adequately perform Stu2’s kinetochore function when only low levels of Stu2 are localized to the kinetochore in the absence of the CTS. The second idea aligns with our previous work, which showed that specific residues within the human ch-TOG basic linker are required for proper chromosome segregation, suggesting that the basic linker contributes to the kinetochore function of the ch-TOG family (Herman et al., 2020). Future work will be required to address these possibilities.

<u>Supplementary discussion point 2</u>: Here we identified a separation of function *stu2* mutant (*stu2^ΔBL^-NLS^SV40^*) that is incapable of polymerizing microtubules in vitro yet is able to support cellular viability. These data indicate that Stu2’s ability to act as a microtubule polymerase is dispensable. However, it is possible that, in cells, Stu2^ΔBL^-NLS^SV40^ can still bind to the microtubule plus-end through an interaction with another microtubule-associated protein (e.g., its known interactors Bik1 or Bim1; Stangier et al., 2018) and subsequently regulate microtubule dynamics. Our data, however, do not support this idea. In vivo, *stu2^ΔBL^-NLS^SV40^* cells exhibit slower anaphase spindle elongation rates, indicating a defect in microtubule polymerization activity. Furthermore, our in vitro microtubule dynamics assay uses whole cell lysates, which include any potential Stu2 binding partners. Our assay also shows that Stu2^ΔBL^-NLS^SV40^ can track the microtubule plus-end, possibly through interaction with another binding partner or via its TOG domains. Despite this localization, Stu2^ΔBL^-NLS^SV40^ fails to properly regulate microtubule dynamics. Supporting our conclusion that microtubule polymerization is not Stu2’s essential function, Geyer et al. demonstrated that although the recombinant Stu2^ΔTOG2^ mutant is deficient as a microtubule polymerase in vitro, it largely sustains cell viability (Geyer et al., 2018).

---

**Figure 1 supplement.**
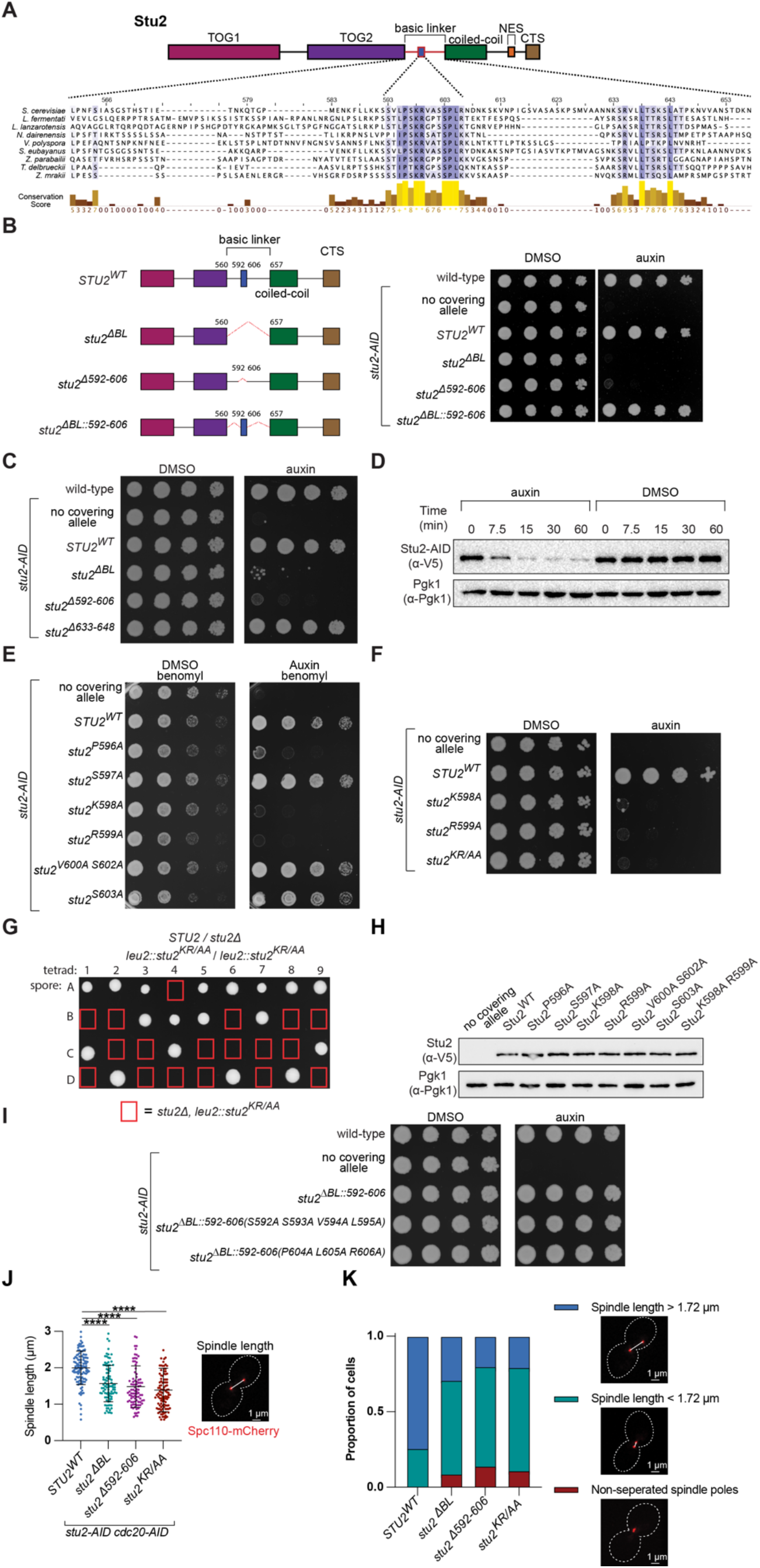
Conservation of the entire basic linker region of Stu2 across 9 fungal species as well as viability and spindle morphology defects of various mutants within Stu2^592-606^. (A) Top: Schematic of Stu2’s domains as in Figure 1A. Bottom: Multiple sequence alignment across 9 different fungal species of Stu2’s entire basic linker region. Note: numbers listed correspond to *S. cerevisiae* amino acid positions. (B) Left: A schematic of different Stu2 mutants tested in Fig. S1B. Right: Cell viability was analyzed in a wild-type strain (M3) as well as *stu2-AID* strains without a covering allele (M619) or ectopically expressing *stu2* alleles (*STU2^WT^*, M622; s*tu2^ΔBL^*, M973; s*tu2^Δ592-606^*, M852; or s*tu2 ^ΔBL::592-606^*, M991) by plating five-fold serial dilutions in the presence of DMSO (left) or auxin (right) to degrade the endogenous Stu2-AID protein. (C) Cell viability was analyzed in a wild-type strain (M3) as well as *stu2-AID* strains without a covering allele (M619) or ectopically expressing *stu2* alleles (*STU2^WT^*, M622; s*tu2^ΔBL^*, M973; s*tu2^Δ592-606^*, M852; or s*tu2^Δ633-648^*, M851) by plating five-fold serial dilutions in the presence of DMSO (left) or auxin (right) to degrade the endogenous Stu2-AID protein. Note: we observed two patches of conservation within the basic linker domain, 592-606 (containing 3 Lys and Arg residues) and 633-648 (containing 4 Lys and Arg residues). Deletion of 592-606 displayed severe growth defects, whereas deletion of 633-648 showed no growth defect. Also see (Stewart et al., 2024). (D) Exponentially growing *stu2-AID-3V5* strain (M608) was split into two cultures, one culture was treated with auxin to degrade *stu2-AID-3V5*, and the other was treated with DMSO. Over the time course of 60 min, samples were harvested from each culture. Protein lysates were subsequently prepared and degradation of Stu2-AID-3V5 proteins was analyzed by immunoblotting. Pgk1 is shown as a loading control. (E) Cell viability was analyzed in *stu2-AID* strains without a covering allele (M619) or ectopically expressing *stu2* alleles (*STU2^WT^*, M622; *stu2^P596A^*, M1290; *stu2^S597A^*, M1299; *stu2^K598A^*, M954; *stu2^R599A^*, M956; *stu2^V600A^ ^S602A^*, M1383; or *stu2^S603A^*, M802) by plating five-fold serial dilutions in the presence of 6.5 µg/mL benomyl with DMSO (left) or auxin (right) to degrade the endogenous Stu2-AID protein. On plates containing benomyl and auxin, cells expressing *stu2^P596A^* display viability defects. (F) Cell viability was analyzed in *stu2-AID* strains without a covering allele (M619) or ectopically expressing *stu2* alleles (*STU2^WT^*, M622; s*tu2^K598A^*, M954; s*tu2^R599A^*, M956; or s*tu2^KR/AA^*, M888) by plating five-fold serial dilutions in the presence of DMSO (left) or auxin (right) to degrade the endogenous Stu2-AID protein. Here we observe that the *stu2^KR/AA^* mutant displays similar viability defects as the two single mutants. (G) Spore viability was analyzed from a sporulated diploid strain (M1265) with a heterozygous deletion of *stu2* at the endogenous *STU2* locus (*STU2*/*stu2Δ*) and homozygous *stu2^KR/AA^*/*stu2^KR/AA^* expressed from the *LEU2* locus. Individual haploid spores were isolated by tetrad dissection and grown on YPD. Each column of four spores is from a single dissected tetrad. Red boxes indicate the location of a spore that contains *stu2Δ* at the endogenous *STU2* locus and *stu2^KR/AA^*at the *LEU2* locus. Lack of colony formation indicates viability defects, consistent with Stu2^K598^ ^R599^ being critical residues for *S. cerevisiae* viability. (H) Exponentially growing *stu2-AID* strains without a covering allele (M619) or ectopically expressing *stu2-3V5* alleles (*STU2^WT^*, M622; *stu2^P596A^*, M1290; *stu2^S597A^*, M1299; s*tu2^K598A^*, M954; *stu2^R599A^*, M956; *stu2^V600A^ ^S602A^*, M1383; *stu2^S603A^*, M802; or *stu2^K598A^ ^R599A^,* M888) were treated with auxin 30 min prior to harvesting. Protein lysates were subsequently prepared and expression of Stu2-3V5 proteins were analyzed by immunoblotting. Pgk1 is shown as a loading control. Note: This is the same blot as in Fig. 1C with the addition of a strain expressing *stu2^K598A^ ^R599A^* (M888). (I) Cell viability was analyzed in a wild-type strain (M3) as well as *stu2-AID* strains without a covering allele (M619) or ectopically expressing *stu2* alleles (*stu2^ΔBL::592-606^,* M991; *stu2^ΔBL::592-^* 606(S592A S593A V594A L595A)_,_ M1519; or stu2*^ΔBL::592-606(P604A L605A R606A)^*^,^ M1518) by plating five-fold serial dilutions in the presence of DMSO (left) or auxin and DMSO (right) to degrade the endogenous Stu2-AID protein. Note: Residues S592, S593, V594, L595, P604, L605, and R606 in the conserved patch (Stu2^592-606^) are not essential for cell viability. (J) Exponentially growing *stu2-AID cdc20-AID* cells ectopically expressing *stu2* alleles (*STU2^WT^*, M1914; *stu2^ΔBL^*, M1918; *stu2^Δ592-606^*, M1921; or *stu2^KR/AA^*, M1922) were treated with auxin for 2 h to degrade Stu2-AID and Cdc20-AID, arresting cells in metaphase. Cells were fixed and the Spc110-mCherry signal was imaged. In cells with bipolar mCherry punctae, the distance was measured between these punctae and is indicated by a data point for each cell. Mean and standard deviation are shown. n = 90–116 cells; P values were determined using an ordinary one-way ANOVA followed by a Tukey’s multiple comparisons test (**** = p<0.0001). (K) Same data as in Fig. S1J quantified to show: (Left) proportion of cells with non-separated spindle pole bodies vs a short spindle (less than the 1^st^ quartile spindle length for cells expressing *STU2^WT^*) vs a normal length spindle (greater than the 1^st^ quartile spindle length for cells expressing *STU2^WT^*) were determined. n = 102-118 cells. (Right) representative cell with varying spindle lengths.

**Figure 2 supplement.**
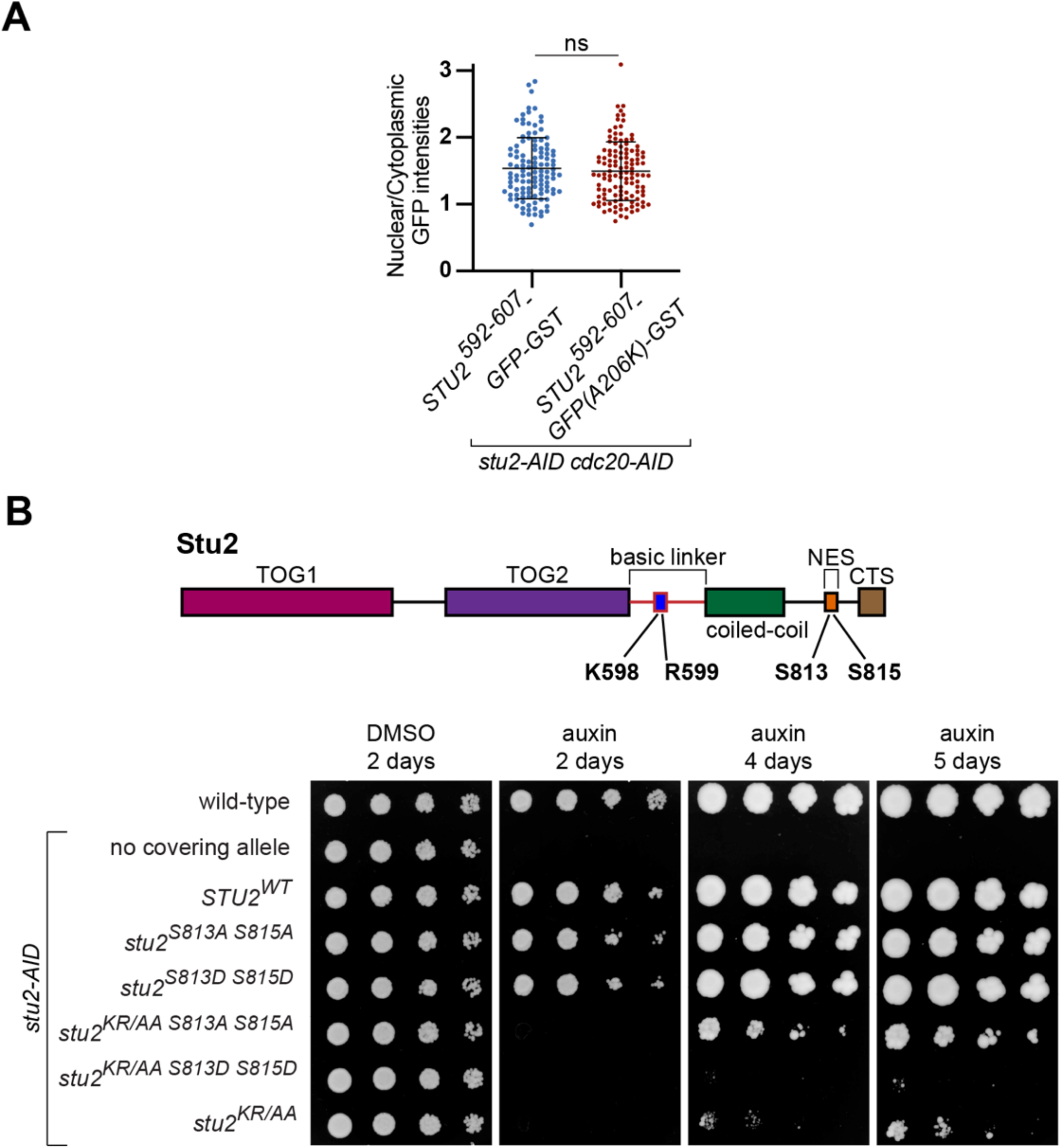
Ratio of nuclear to cytoplasmic GFP signal intensity in Stu2^592-607^-GFP-GST with a copy of GFP that is unable to dimerize and modulating nuclear levels of Stu2^KR/AA^ affects viability. (A) Exponentially growing *stu2-AID cdc20-AID* cells ectopically expressing *GFP-GST* fusion constructs (*stu2^592-607^-GFP-GST*, M2392; or *stu2^592-607^-GFP(A206K)-GST*, M2480) were treated with auxin for 2 h to degrade Stu2-AID and arrest cells in metaphase. Cells were fixed and Nup2-mKATE and GFP signals were imaged and the ratio of nuclear to cytoplasmic GFP signal intensities was quantified. Each data point represents this ratio for a single cell. Mean and standard deviation are shown. n=116-122 cells; P values were determined using a two-tailed unpaired *t* test. Here we find that using a copy of GFP that is unable to homodimerize, as shown in (Zacharias et al., 2002), does not alter the nuclear to cytoplasmic GFP ratio of Stu2^592-607^- GFP-GST, suggesting that the conserved patch (Stu2^592-607^) does not need to be a dimer for nuclear localization function. (B) Top: Schematic of Stu2 domains indicating mutated residues in the experimental strains used in Fig S2B. Bottom: Cell viability was analyzed in a wild-type strain (M3) as well as *stu2- AID* strains without a covering allele (M619) or ectopically expressing *stu2* alleles (*STU2^WT^*, M2103; stu2S8*^13A S815A^*, M2248; s*tu2^S813D S815D^*_,_ M2249; s*tu2^KR/AA S813A S815A^*_,_ M2250; s*tu2^KR/AA S813D S815D^*, M2251; or s*tu2^KR/AA^*, M2225) by plating five-fold serial dilutions in the presence of DMSO (left) or auxin (right) to degrade the endogenous Stu2-AID protein. Auxin plate was imaged over the course of 2-5 days. Note: Stu2’s nuclear export signal (NES) is regulated via Sch9- dependent phosphorylation of Stu2^S813^ ^S815^, thereby promoting Stu2’s nuclear export (van der Vaart et al., 2017). When we combined *stu2^KR/AA^*with phosphorylation mimicking mutants (S813D S815D), thus decreasing the nuclear pool of Stu2 by promoting nuclear export, we observe an exacerbation of the viability defects seen in *stu2^KR/AA^* expressing cells. Conversely, when we increased the nuclear pool of Stu2^KR/AA^ by combining with the phosphorylation deficient mutations (S813A S815A), thus inhibiting Stu2 nuclear export, we observe a partial rescue of the *stu2^KR/AA^* viability phenotype.

**Figure 3 supplement.**
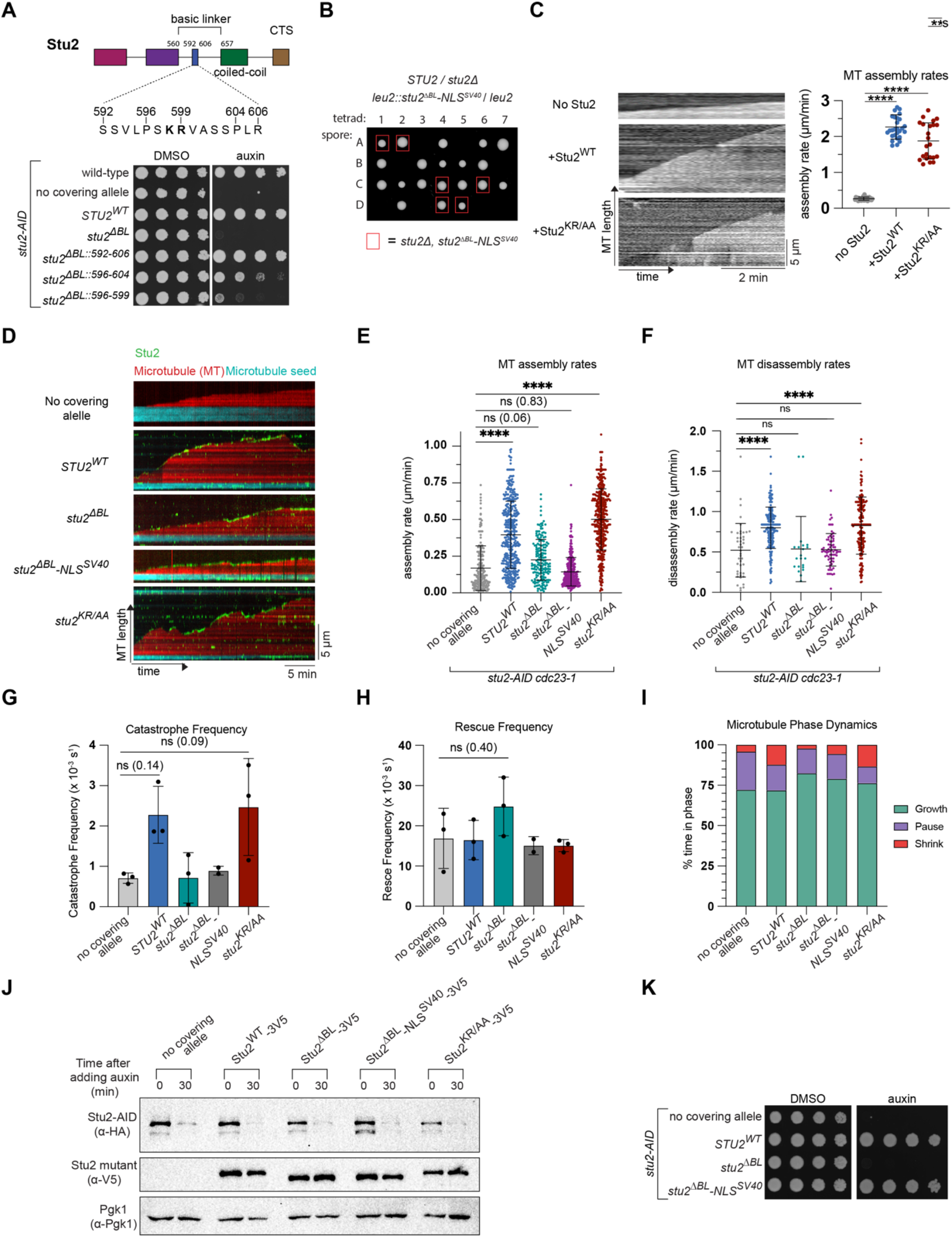
The basic linker domain is required for Stu2’s ability to stimulate microtubule assembly and disassembly in vitro but is dispensable for cell viability as long as Stu2 is localized to the nucleus. (A) Cell viability was analyzed in a wild-type strain (M3) as well as *stu2-AID* strains without a covering allele (M619) or ectopically expressing *stu2* alleles (*STU2^WT^*, M622; *stu2^ΔBL^*, M973; *stu2^ΔBL::592-606^*, M991; *stu2^ΔBL::596-604^*, M1710; *stu2^ΔBL::596-599^*, M1301) by plating five-fold serial dilutions in the presence of DMSO (left) or auxin (right) to degrade the endogenous Stu2-AID protein. (B) Spore viability was analyzed from a sporulated diploid strain (M5676) with a heterozygous deletion of *stu2* at the endogenous *STU2* locus (*STU2*/*stu2Δ*) and a heterozygous insertion of *stu2^ΔBL^-NLS^SV40^* at the *leu2* locus (*stu2^ΔBL^-NLS^SV40^*/*leu2*). Individual haploid spores were isolated by tetrad dissection and grown on YPD. Each column of four spores is from a single dissected tetrad. Red boxes indicate the location of a spore that contains *stu2Δ* at the endogenous *stu2* locus and *stu2^ΔBL^-NLS^SV40^*at the *leu2* locus. Viability of these spores indicates that Stu2^ΔBL^- NLS^SV40^ is capable of supporting *S. cerevisiae* viability when it is the only copy available in the cell, consistent with the idea that driving nuclear localization is the only essential function carried out by the basic linker. (C) Left: Representative kymographs for αβ-tubulin alone, +Stu2^WT^, or +Stu2^KR/AA^. Microtubule dynamics were measured by time-lapse DIC microscopy. Right: Quantification of MT growth rates with yeast αβ-tubulin (0.8 μM) without Stu2 or with 250 nM Stu2^WT^ or Stu2^KR/AA^, as indicated. Average growth rates for αβ-tubulin, αβ-tubulin+Stu2^WT^ and αβ-tubulin+Stu2^KR/AA^ were 0.267 ± 0.045 µm/min (n=26), 2.269 ± 0.333 µm/min (n= 28), and 1.879 ± 0.493 µm/min (n=23), respectively. Mean and standard deviation are shown. P values were determined using an ordinary one-way ANOVA followed by a Tukey’s multiple comparisons test (**** = p<0.0001). (D) Exponentially growing *GFP-TUB1 stu2-AID cdc23-1* cells with no covering allele (M2166) or ectopically expressing *stu2-Halo* (*STU2^WT^*, M2231; *stu2^ΔBL^*, M2360; *stu2^ΔBL^-NLS^SV40^*, M3516, or *stu2^KR/AA^*, M2232) were shifted to a non-permissive temperature for 3 h to arrest cells in metaphase and treated with auxin 30 min before harvesting to degrade Stu2-AID. Whole cell lysate was incubated with microtubule seeds and imaged over time with TIRF microscopy. Representative kymograph images for individual microtubule extensions. These kymographs show microtubule extension (red) from the microtubule seed (cyan) and Stu2^mut^-Halo (green) over time. Note: this is the same data as in Fig. 3D with the added strain *stu2^KR/AA^* (M2232). (E) Microtubule assembly rates were calculated from the kymographs (representative kymographs in Fig. S3D). Each data point represents an independent microtubule assembly event. Mean and standard deviation are shown. n=182-408 growth events; P values were determined using an ordinary one-way ANOVA followed by a Tukey’s multiple comparisons test (**** = p<0.0001). (F) Microtubule disassembly rates were calculated from the kymographs (representative kymographs in Fig. S3D). Each data point represents an independent microtubule disassembly event. Mean and standard deviation are shown. n=21-191 shrinkage events; P values were determined using an ordinary one-way ANOVA followed by a Tukey’s multiple comparisons test (**** = p<0.0001). (G) Microtubule catastrophe frequencies were calculated from the kymographs (representative kymographs in Fig. S3D). Each data point represents the average catastrophe rate from all kymographs of one experimental replicate. Mean and standard deviation are shown. n=2-3 experimental replicates; P values were determined using an ordinary one-way ANOVA followed by a Tukey’s multiple comparisons test. Note: catastrophe frequency increases by approximately 3.2-fold from no covering allele to *STU2^WT^*. While this difference is not statistically different in Tukey’s multiple comparisons test, likely because of the low number of data points, it suggests that Stu2 is a microtubule catastrophe promoting agent and that this activity depends on the basic linker. (H) Microtubule rescue frequencies were calculated from the kymographs (representative kymographs in Fig. S3D). Each data point represents the average catastrophe rate from all kymographs of one experimental replicate. Mean and standard deviation are shown. n=2-3 experimental replicates; P values were determined using an ordinary one-way ANOVA followed by a Tukey’s multiple comparisons test. (I) The average percent time that a microtubule was in a state of growth versus shrinkage versus paused are indicated (representative kymographs in Fig. S3D). (J) Exponentially growing *GFP-TUB1 stu2-AID cdc23-1* cells with no covering allele (M2166) or ectopically expressing *stu2-Halo* (*STU2^WT^*, M2231; *stu2^ΔBL^*, M2360; *stu2^ΔBL^-NLS^SV40^*, M3516, or *stu2^KR/AA^*, M2232) were shifted to a non-permissive temperature for 3 h to arrest cells in metaphase and treated with auxin 30 min before harvesting to degrade Stu2-AID. Protein lysates were prepared at 2.5 h (before adding auxin) and at 3 h (after adding auxin) and expression of Stu2-AID-HA and Stu2-3V5 proteins were analyzed by immunoblotting. Pgk1 is shown as a loading control. Note: these are the same strains treated similarly as in Fig. S3C,. The lysates prepared for the western blot are separate from those used in the microtubule growth dynamics assay. (K) Cell viability was analyzed in *stu2-AID* strains without a covering allele (M2166) or ectopically expressing *stu2-Halo-3V5* alleles (*STU2^WT^*, M2231; *stu2^ΔBL^,* M2360; or *stu2^ΔBL^- NLS^SV40^,* M3516) by plating five-fold serial dilutions in the presence of DMSO (left) or auxin (right) to degrade the endogenous Stu2-AID protein. Note: These are the strains used for the whole cell lysate assay.

**Figure 4 supplement.**
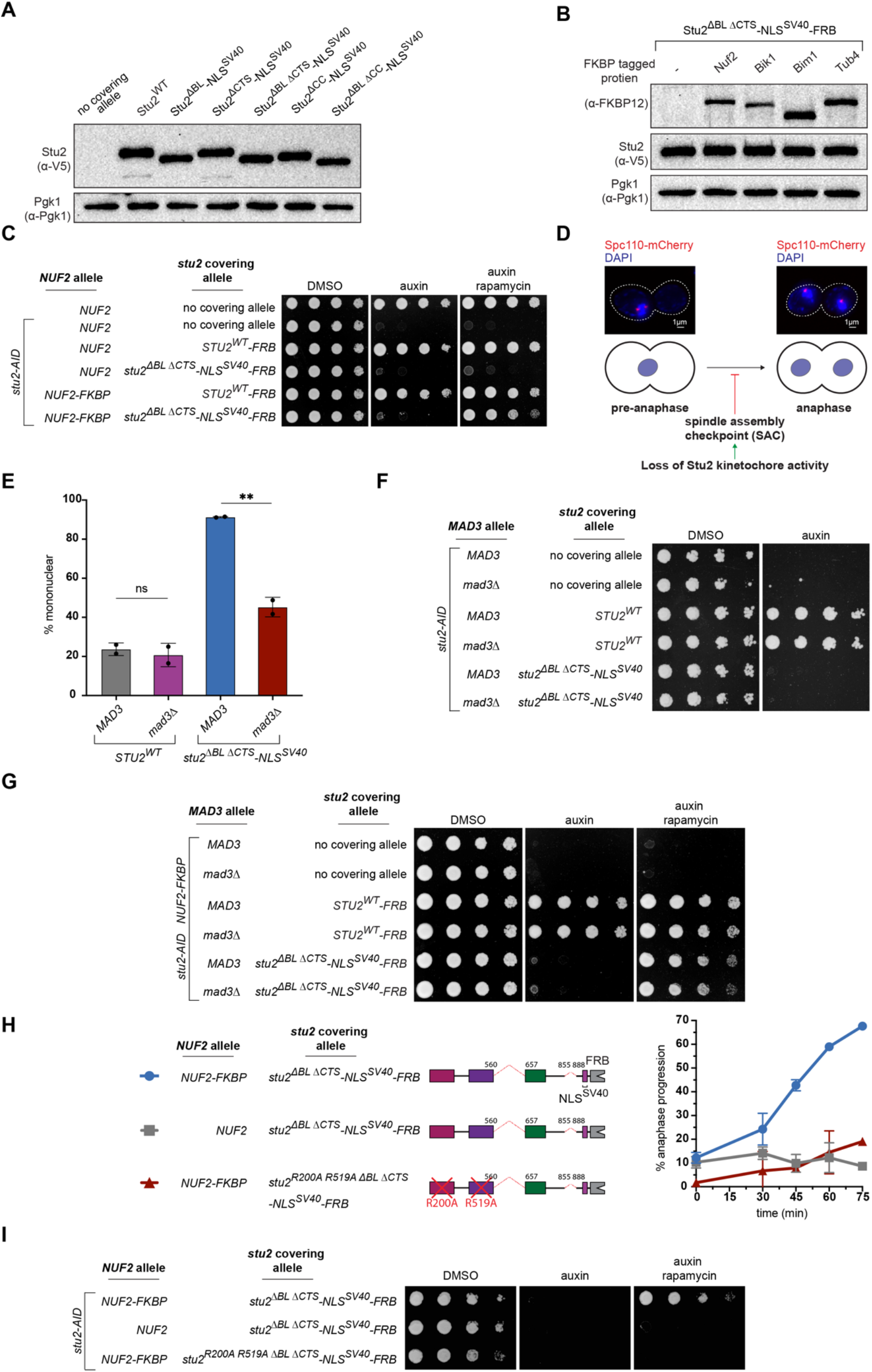
Stu2’s kinetochore function is required for cell viability independently of the SAC activity. (A) Samples of exponentially growing *stu2-AID* strains without a covering allele (M619) or ectopically expressing *stu2-3V5* alleles (*STU2^WT^*, M622; *stu2^ΔBL^-NLS^SV40^*, M3722; *stu2^ΔCTS^- NLS^SV40^*, M3942; *stu2^ΔBL^ ^ΔCTS^-NLS^SV40^*, M3727; *stu2^ΔCC^-NLS^SV40^*, M3940; or *stu2^ΔCC^ ^ΔCTS^-NLS^SV40^*, M3729) were harvested. Protein lysates were subsequently prepared, and expression of Stu2- 3V5 proteins was analyzed by immunoblotting. Pgk1 is shown as a loading control. (B) Samples of exponentially growing *stu2-AID TOR1-1 fpr1Δ* strains with *stu2^ΔBL^ ^ΔCTS^-NLS^SV40^- FRB* covering allele and without any FKBP tagged gene (M4515) or endogenously expressing FKBP tagged genes (*NUF2*, M4366; *BIK1*, M4369; *BIM1*, M4371; or *TUB4*, M4367) were harvested. Protein lysates were subsequently prepared and expression of FKBP tagged protein and stu2^ΔBL, ΔCTS^-NLS^SV40^-FRB-3V5 protein were analyzed by immunoblotting. Pgk1 is shown as a loading control. (C) Cell viability was analyzed in a *TOR1-1 fpr1Δ* strain that is otherwise wild-type (M1375) and *TOR1-1 fpr1Δ stu2-AID* strains with *NUF2^WT^* and without a covering allele (M1477) or ectopically expressing *stu2* alleles (*STU2^WT^-FRB*, M4514; or *stu2^ΔBL^ ^ΔCTS^-NLS^SV40^-FRB*, M4515) or with *NUF2-FKBP* and ectopically expressing *stu2* alleles (*STU2 ^WT^-FRB*, M4491; or *stu2^ΔBLΔCTS^-NLS^SV40^-FRB*, M4366) by plating five-fold serial dilutions in the presence of DMSO (left), auxin (middle) to degrade the endogenous Stu2-AID protein, or auxin and rapamycin (right) to induce the dimerization of FRB and FKBP and degrade the endogenous Stu2-AID protein. (D) Bottom: a schematic of large-budded mononucleate cells and the resulting binucleate cells that result from anaphase onset. Loss of Stu2’s activity leads to activation of the spindle assembly checkpoint (SAC), arresting the cells prior to anaphase. Top: Representative cells with DNA stained with DAPI (blue) indicating the nucleus and the spindle pole bodies (SPBs) marked with Spc110-mCherry (red). (E) Exponentially growing *SPC110-mCherry stu2-AID* cells expressing an ectopic copy of Stu2 either *with MAD3 or mad3Δ*, to have either an active SAC or an inactive SAC, were treated with auxin for 2 h to degrade the endogenous Stu2-AID protein and to allow cells enough time to arrest in mitosis if the SAC is activated (*STU2^WT^ MAD3^WT^*, M4445; *STU2^WT^ mad3Δ*, M4497; *stu2^ΔBL^ ^ΔCTS^-NLS^SV40^ MAD3^WT^*, M4509; or *stu2^ΔBL^ ^ΔCTS^-NLS^SV40^ mad3Δ*, M4505). Cells were fixed and Spc110-mCherry and DAPI signals were imaged. Percentage of mononucleate large-budded cells was quantified out of the total population of large-budded cells, non-large-budded cells were excluded from analysis. Each data point is a biological replicate, n=109-130 large-budded cells for each replicate. Mean and standard deviation are shown. P values were determined using a two-tailed unpaired *t* test (** = p<0.01). (F) Cell viability was analyzed in *stu2-AID* strains without a covering allele either *with MAD3* (M2398) or *mad3Δ* (M4503), to have either an active SAC or an inactive SAC or with *stu2* covering allele (*STU2^WT^ MAD3^WT^*, M4445; *STU2^WT^ mad3Δ*, M4497; *stu2^ΔBL^ ^ΔCTS^-NLS^SV40^MAD3^WT^*, M4509; or *stu2^ΔBL^ ^ΔCTS^-NLS^SV40^ mad3Δ*, M4505) by plating five-fold serial dilutions in the presence of DMSO (left), or auxin (right) to degrade the endogenous Stu2-AID protein. (G) Cell viability was analyzed in *stu2-AID NUF2-FKBP TOR1-1 fpr1Δ* strains without a covering allele either with *MAD3* (M1944) or *mad3Δ* (M1986), to have either an active SAC or an inactive SAC or with a *stu2* covering allele (*STU2^WT^-FRB MAD3^WT^*, M4510; *STU2^WT^-FRB mad3Δ*, M4511; *stu2^ΔBL^ ^ΔCTS^-NLS^SV40^-FRB MAD3^WT^*, M4512; or *stu2^ΔBL^ ^ΔCTS^-NLS^SV40^-FRB mad3Δ*, M4513) by plating five-fold serial dilutions in the presence of DMSO (left), auxin (middle) to degrade the endogenous Stu2-AID protein, or auxin and rapamycin (right) to induce the dimerization of FRB and FKBP and degrade the endogenous Stu2-AID protein. (H) Left: A schematic of mutants used in the experiment. Right: Exponentially growing *SPC110- mCherry stu2-AID TOR1-1 fpr1Δ* cells with (*stu2^ΔBL^ ^ΔCTS^-NLS^SV40^-FRP NUF2-FKBP*, M4318; *stu2^ΔBL^ ^ΔCTS^-NLS^SV40^-FRP NUF2^WT^*, M4354; or *stu2^R200A^ ^R519A^ ^ΔBL, ΔCTS^-NLS^SV40^-FRP NUF2-FKBP*, M4576). were treated with auxin for 2 h to degrade the endogenous Stu2-AID protein and exhibit a pre-anaphase arrest. After 2 h rapamycin was added (T0) to induce FRB/FKBP dimerization and samples were collected and fixed over a time course of 75 min. Spc110- mCherry and DAPI signals were imaged. Percentage of large-budded cells that progressed through anaphase out of the total population of large-budded cells (non-large-budded cells were excluded from analysis) was quantified for each time point. Data represents two biological replicates, n=100-145 large-budded cells for each time point and each replicate. Mean and standard deviation of both replicates are shown at each time point. Same data as in Fig. 4C with the added *stu2^R200A^ ^R519A^ ^ΔBL, ΔCTS^-NLS^SV40^-FRP* mutant. (I) Cell viability was analyzed in *stu2-AID TOR1-1 fpr1Δ* strains with a *stu2^ΔBL^ ^ΔCTS^-NLS^SV40^-FRB* covering allele and endogenously expressed *NUF2-FKBP* (M4318) or without any FKBP tagged gene (M4354) or with a *stu2^R200A^ ^R519A^ ^ΔBL^ ^ΔCTS^-NLS^SV40^-FRB* covering allele and endogenously expressed *NUF2-FKBP* (M4576) by plating five-fold serial dilutions in the presence of DMSO (left), auxin (middle) to degrade the endogenous Stu2-AID protein, or auxin and rapamycin (right) to induce the dimerization of FRB and FKBP and degrade the endogenous Stu2-AID protein.

**Figure 5 supplement.**
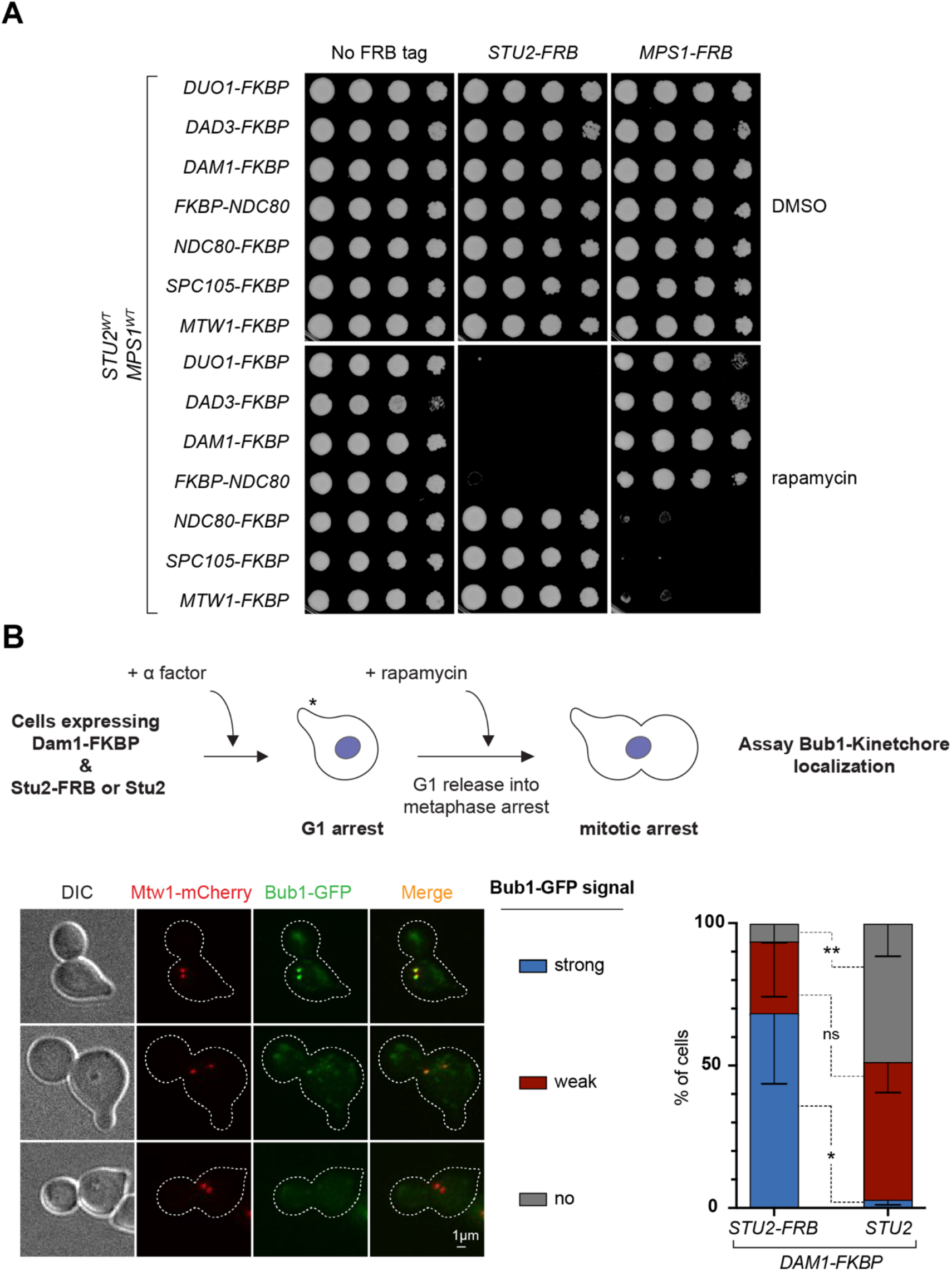
Stu2’s precise kinetochore position is critical for function. (A) Cell viability was analyzed by plating five-fold serial dilutions in the presence of DMSO (top) or rapamycin (bottom) to induce the dimerization of FRB and FKBP. Strains contained endogenous *STU2^WT^*and *MPS1^WT^* with *TOR1-1 fpr1Δ* and an FKBP tagged protein (*DUO1- FKBP*, M5519; *DAD3-FKBP*, M5516; *DAM1-FKBP*, M4784; *FKBP-NDC80*, M5510; *NDC80- FKBP*, M1427; *SPC105-FKBP*, M4613; or *MTW1-FKBP*, M4614) in the absence of an FRB tagged protein (Left), or also containing an ectopic *STU2-FRB* (Middle; with an FKBP tagged protein: *DUO1-FKBP*, M5673; *DAD3-FKBP*, M5643; *DAM1-FKBP*, M5655; *FKBP-NDC80*, M5637; *NDC80-FKBP*, M5631; *SPC105-FKBP*, M5634; or *MTW1-FKBP*, M5646), or with ectopic *MPS1-FRB* (Right; with an FKBP tagged protein: *DUO1-FKBP*, M5675; *DAD3-FKBP*, M5645; *DAM1-FKBP*, M5657; *FKBP-NDC80*, M5639; *NDC80-FKBP*, M5633; *SPC105-FKBP*, M5636; or *MTW1-FKBP*, M5648). Same data as in Fig. 5B with the added DMSO control plates. (B) Top: A schematic of the experimental design. Exponentially growing *DAM1-FBKP BUB1- GFP MTW1-mCherry pMET-cdc20 TOR1-1 fpr1Δ* cells expressing an ectopic copy of *STU2^WT^*(M5577) or *STU2^WT^-FRB* (M5521) were treated with α factor (mating pheromone) for 3 h to synchronize cells in G1 in synthetic media lacking L-methionine. The asterisk denotes the mating projection (a shmoo) induced by α factor. After 3 h, cells were released from G1 arrest into YPAD media + L-methionine to arrest in metaphase (as L-methionine represses the Met promotor of *pMET-cdc20*) + rapamycin. After 2 h, samples were collected and fixed. Bub1-GFP and Mtw1-mCherry signals were imaged. Percentage of large-budded cells with a strong signal of Bub1-GFP colocalized to Mtw1-mCherry signal, weak Bub1-GFP signal and no Bub1-GFP signal was quantified. A cut off value of Bub1-GFP signal was used to differentiate strong vs weak signals. Bottom-left: Representative images of different Bub1-GFP signals. Bottom-right: Quantification of percentage of cells with different Bub1-GFP signals. The graphs represent three biological replicates, n=71-126 for each replicate. Mean and standard deviation are shown. P values were determined using a two-tailed unpaired *t* test (* = p<0.05 and ** = p<0.01).

